# RNA polymerase II assembly and mRNA decay regulation are mediated and interconnected *via* CTD Ser5P phosphatase Rtr1 in *Saccharomyces cerevisiae*

**DOI:** 10.1101/2020.12.23.424112

**Authors:** A.I. Garrido-Godino, A. Cuevas-Bermúdez, F. Gutiérrez-Santiago, M.C. Mota-Trujillo, F. Navarro

## Abstract

Rtr1 is an RNA pol II CTD-phosphatase that influences gene expression by acting during the transition from transcription initiation to elongation, and during transcription termination. Rtr1 has been proposed as an RNA pol II import factor in RNA pol II biogenesis, and participating in mRNA decay by autoregulating the turnover of its own mRNA. In addition, the interaction of Rtr1 with RNA pol II depends on the phosphorylation state of CTD, which also influences Rpb4/7 dissociation during transcription. In this work, we demonstrate that Rtr1 acts in RNA pol II assembly, likely in a final cytoplasmic RNA pol II biogenesis step, and mediates the Rpb4 association with the rest of the enzyme, However, we do not rule out discard a role in the Rpb4 association with RNA pol II in the nucleus. This role of Rtr1 interplays RNA pol II biogenesis and mRNA decay regulation. In fact, *RTR1* deletion alters RNA pol II assembly and leads to the chromatin association of RNA pol II lacking Rpb4, in addition to whole RNA pol II, decreasing mRNA-Rpb4 imprinting and, consequently, increasing mRNA stability. Notably, the *RPB5* overexpression that overcomes RNA pol II assembly and the defect in Rpb4 binding to chromatin-associated RNA pol II partially suppresses the mRNA stability defect of *rtr1Δ* cells. Our data also indicate that Rtr1 mediates mRNA decay regulation more broadly than previously proposed in cooperation with Rpb4 and Dhh1. Interestingly, these data include new layers in the crosstalk between mRNA synthesis and decay.

## INTRODUCTION

Transcription is the most widely studied step in gene expression. In most eukaryotes, transcription is carried out by three heteromultimeric RNA polymerases named RNA pol I, RNA pol II and RNA pol III that synthesize different types of RNAs [1]. In plants, RNA pol IV and V also exist that transcribe regulatory ncRNAs [2,3]. Although the eukaryotic RNA polymerases structure [4,5,6,7,8,9] and many aspects of transcription process [5,10,11,12,13] have been extensively studied, the global assembly of RNA polymerases has barely been explored.

Biogenesis of eukaryotic multisubunit RNA polymerases has been proposed to be similar to the bacterial type by assuming the formation of three subassembly complexes: the Rpb1 (composed of Rpb1, Rpb5, Rpb6 and Rpb8), Rpb2 (consisting of Rpb2 and Rpb9) and Rpb3 (comprising Rpb3, Rpb10, Rpb11 and Rpb12) subassemblies. These subcomplexes interact to form the complete enzyme prior to its nuclear import [14]. Some factors have been described to be involved in the assembly of RNA pol II in human cells, such as the HSP90 cochaperone or the R2TP/prefoldin-like complex [15]. Similarly, small GTPases GPN1, GPN2 and GPN3 in human [16] and their yeast counterparts, as well as yeast Rtp1 or Rba50, have been proposed to participate in RNA Pol II assembly [17,18,19,20]. However, very little is known about the factors that mediate the assembly of RNA pols I and III or the assembly of the three enzymes. Yeast small GTPases Gpn2 and Gpn3, which also participate in RNA pol II assembly, mediate the assembly of RNA pol III [19], while yeast protein Rbs1 has been proposed to be involved in RNA pol III biogenesis [21]. Only prefoldin-like Bud27 has been described as a factor that mediates the assembly of the three RNA pols in a process that depends on the common subunit to the three RNA pols, Rpb5 [22,23]. Bud27 mediates the association of Rpb5 and Rpb6 with the rest of the complex prior to its nuclear import [22]. The Bud27 human counterpart, URI, is described to form part of the HSP90/R2TP-prefoldin-like complex, which participates in the cytoplasmic assembly of RNA pol II [15,16,22,24]. In addition to RNA pol II assembly factors, other proteins have been shown to participate in RNA pol II transport to the nucleus in *Saccharomyces cerevisiae*, such as Iwr1 [25,26], Rtp1 [18] and Rtr1 [26]. The nuclear import of RNA pol II in *S. cerevisiae* occurs mainly through the Iwr1-dependent process [25,26]. However, the largest RNA pol II subunits can also be imported to the nucleus by an Iwr1-independent pathway with both the participation of Rtr1 and Rtp1 and the action of microtubules [26]. This Iwr1-independent pathway also seems to operate for the smallest RNA pol II subunits, which can enter the nucleus by diffusion [26].

Rtr1 (for “regulator of transcription” 1) and its human ortholog RPAP2 (for “RNAPII-associated polypeptide”) were initially described as RNA polymerase II interactors [27,28]. Rtr1 is an S5-P CTD phosphatase with a proposed role in the transition from transcription initiation to elongation *in vivo* [29,30]. Furthermore, Rtr1 can dephosphorylate Tyr1-P CTD *in vitro*, which suggests a role in transcription elongation and termination phases [31]. However, the phosphatase activity of Rtr1 is controversial as Rtr1 seems to either lack the typical active site in *Kluyveromyces lactis* [32] or possess an active site for atypical phosphatase in *S. cerevisiae* [33]. In addition, Rtr1 has been proposed to associate with RNA pol II during transcription, which also seems the case of other elongation factors, depending on the CTD phosphorylation state. This interaction influences the Rpb4/7 dissociation with the rest of the enzyme [34]. Finally, Rtr1 and RPAP2 are implicated in the transcription of non-coding RNAs [35,36,37].

Despite the effect of Rtr1 in transcription, some studies have proposed a role for Rtr1 and its human ortholog RPAP2 in the biogenesis of RNA pol II by acting as transport factors to allow RNA pol II shuttling from the cytoplasm to the nucleus [26,38]. Accordingly, the silencing of RPAP2 or the deletion of *RTR1* causes the accumulation of Rpb1 and Rpb2 in the cytoplasm, the two largest RNA pol II subunits, whereas smaller subunits can reach the nuclear space. Rtr1 localizes in the cytoplasm of cells under both normal growth conditions and at high temperatures [27,39,40], and shuttles between the nucleus and cytoplasm in an Xpo1-dependent manner [27]. Similarly, human RPAP2 can be found in both the nucleus and cytoplasm given its nuclear retention domain at its N-terminal region, as well as its cytoplasmic retention domain located in the C-terminal part of the protein [38]. Interactions between Rtr1 and RPAP2 with RNA pol II are controversial. Some authors propose an interaction between Rtr1 and active phosphorylated RNA pol II [27,29,37,41], while others posit that the efficient RNA pol II/RPAP2 interaction does not require the CTD domain of the enzyme in either *in vitro* or *in vivo* [38]. RPAP2 interacts with RNA pol II *in vitro* through its nuclear retention domain [38], likely via C-terminal 460 amino acids, and directly binds the Rpb6 subunit of the enzyme [42]. Furthermore, RNA pol II purification using Rtr1-TAP as bait results in the isolation of a 10-subunit enzyme with the relative depletion of heterodimer Rpb4/7 [34].

In line with a role of Rtr1 as an RNA pol II transporting factor, an interaction between Rtr1 and the nucleocytoplasmic transport protein Ran has been described [43]. Furthermore in Rtr1 purification experiments, interactions with GTPases Gpn3 and Npa3 (the yeast homologue of GPN1) have been detected [34,41], where Npa3 being implicated in RNA pol II nuclear import regulation [20]. In humans, GPN1 interacts with the cytoplasmic retention domain of RPAP2 [28,38] and its silencing results in RPAP2 nuclear retention [38]. RPAP2 has been suggested to return to the cytoplasm in association with GPN1 in a CRM1-dependent process [38].

*RTR1* has a paralogue in *S. cerevisiae* called *RTR2* [27]. While *RTR1* deletion causes Rpb1 accumulation in the cytoplasm, *RTR2* deletion does not provoke Rpb1 mislocalization. However, the *rtr1Δ rtr2Δ* double mutant exacerbates *rtr1Δ* Rpb1 cytoplasmic accumulation, which indicates that Rtr1 and Rtr2 play redundant roles in Rpb1 import [26].

A recent study proposed that Rtr1 plays a role in mRNA stability. This work showed that Rtr1 autoregulates the stability of its own mRNA though its interaction with the corresponding mRNA on a degradation pathway that involves 5’-3’ DExD/H-box RNA helicase Dhh1 and 3’-5’ exonucleases Rex2p and Rex3p [44]. This work opens a way to Rtr1, as transcriptional machinery, to influence the mRNA life cycle from transcription to mRNA decay. Some authors define gene expression as a circular process during which the synthesis and degradation of mRNAs are coordinated in a process called the crosstalk of mRNA, which extend the classic view of the Central Dogma of the Molecular Biology [45,46,47,48,49,50,51]. During this process, mRNA decay machinery may influence transcription and transcription machinery might regulate mRNA fate [46,47,50,51,52]. RNA pol II impacts mRNA stability through its Rpb4 subunit, which binds mRNA during transcription and accompanies it throughout its life cycle by regulating processes like export, translation and decay [48,53,54,55,56]. In fact alterations of RNA pol II assembly, which leads to the dissociation of Rpb4 from the rest of the complex, increase mRNA stability [48,55].

Although Rtr1 can regulate different gene expression steps from RNA pol II biogenesis to transcription and mRNA stability, the specific role of Rtr1 in both assembly and mRNA decay regulation is poorly understood. In this work we demonstrate that Rtr1 acts in RNA pol II assembly to likely favour the correct formation of the Rpb1 subassembly complex and its association, probably in a final step during cytoplasmic RNA pol II biogenesis. Our data also point out that Rtr1 is necessary to mediate the Rpb4 association with the rest of the enzyme (and likely of the Rpb4/7 subassembly complex) in the cytoplasm, although we cannot rule out that this process also occurs in the nucleus. *rtr1Δ* provokes RNA pol II assembly defects that lead to chromatin-associated RNA pol II lacking the Rpb4 subunit, which affects Rpb4-imprinted mRNA and increases mRNA stability. In addition, our data demonstrate that Rtr1 does not play a major role in mediating Rpb4 dissociation from the rest of the enzyme during the transcriptional process. We demonstrate that overcoming the RNA pol II assembly defects of *rtr1Δ* cells by *RPB5* overexpression suffices to restore the decrease in Rpb4 association with the chromatin-associated RNA pol II and, consequently, to partially suppress the mRNA stability alteration. Our data also suggests a specific role of Rtr1 in mRNA decay regulation. Finally, our data also point out a coordinated cooperation among Rtr1, Rpb4 and Dhh1 to modulate mRNA imprinting and mRNA decay.

## RESULTS

### Correct RNA polymerase II assembly is mediated by Rtr1

CTD Ser5P phosphatase Rtr1 and its human homologue RPAP2 have been described as RNA pol II interactors [27,28,29,57], with a proposed role as RNA pol II CTD Ser5 phosphatases [29,30,31,37]. Some laboratories have described a role for Rtr1 and RPAP2 in the biogenesis of RNA pol II by facilitating its transport from the cytoplasm to the nucleus because the deletion of both *RTR1* and silencing of RPAP2 leads to the cytoplasmic accumulation of the largest RNA pol II subunit, Rpb1 [26,38]. However, the possibility of Rtr1 and RPAP2 acting as assembly factors of RNA pol II, which would lead their inactivation to both the disruption of RNA pol II biogenesis and the appearance of RNA pol II subcomplexes, has scarcely been explored.

In an attempt to decipher whether Rtr1 may be involved in the assembly of RNA pol II, we performed protein immunoprecipitation with an anti-Rpb3 antibody against the Rpb3 subunit of RNA pol II in an *rtr1Δ* mutant and its isogenic wild-type strain. Then we analysed the Rpb3/Rpb1 ratio by western blot with an anti-Rpb1 antibody against the largest subunit of the RNA pol II amino-terminal domain (amino acids 1 to 80) to avoid interference with Rpb1 phosphorylation. As shown in Figure 1A and S1A, lack of Rtr1 did not significantly alter the Rpb3/Rpb1 ratio. Similar results were obtained when a polyclonal antibody against the C-terminal domain (CTD) of Rpb1 [58] was used (Figure 1A and S1A). No significant differences in the Rpb3/Rpb1 ratio were observed in either the RNA pol II pull-down experiments (TAP purification) from the *rtr1Δ* mutant and its isogenic wild-type strains, both containing a functional TAP-tagged version of Rpb2, or the Rpb1/Rpb2 or Rpb3/Rpb2 ratios (not shown).

**Figure 1:**
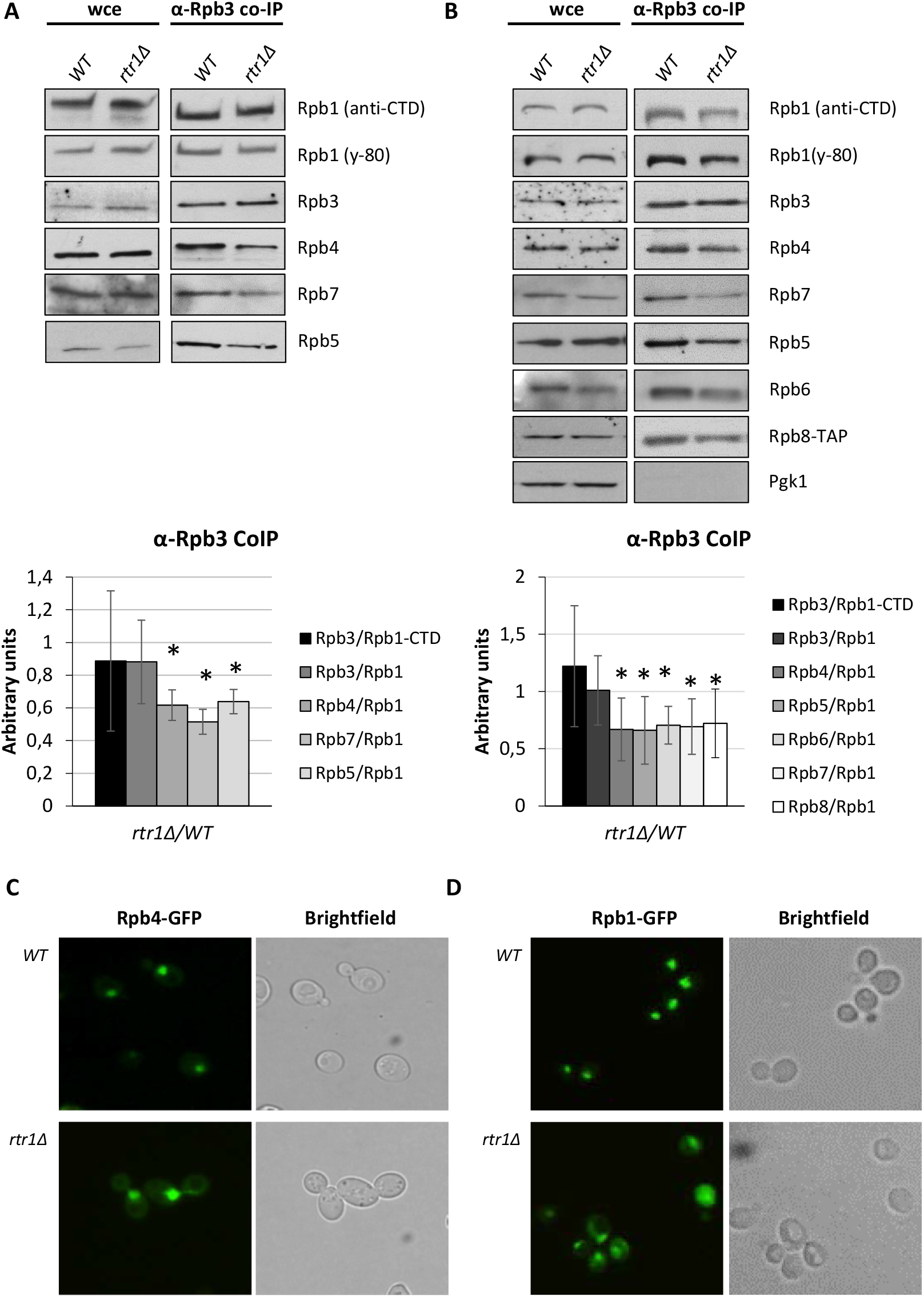
*RTR1* deletion affects the assembly of RNA pol II. **A**) The RNA pol II immunoprecipitated from the wild-type and *rtr1Δ* mutant strains using an anti-Rpb3 antibody and grown in YPD medium at 30°C. The upper panel shows the western blot analysis of subunits Rpb1 (anti-CTD and y-80 antibodies), Rpb3 Rpb4, Rpb7 and Rpb5 of RNA pol II in whole cell extracts, and in the immunoprecipitated samples. Lower panel: quantification of western blot showing the *rtr1Δ* mutant/wild-type strains ratio for each subunit *vs*. Rpb1 (y-80). *p<0.05 (t-test). Pgk1 was tested as a negative control in the RNA pol II purified samples. **B**) Rpb3 immunoprecipitation of RNA pol II using the wild-type and *rtr1Δ* strains containing an Rpb8-TAP tag and grown in YPD medium at 30°C. The western blot of Rpb1 (anti-CTD and y-80 antibodies), Rpb3, Rpb4, Rpb7, Rpb5, Rpb6, and Rpb8 from whole cell crude extracts and immunoprecipitated samples are shown (upper panel). Lower panel: quantifications of western blot showing the *rtr1Δ* mutant/wild-type strains ratio for each subunit *vs*. Rpb1 (y-80). Pgk1 was tested as a negative control in the RNA pol II purified samples. Graphs represent the median and SD of at least three independent biological replicates. **C**) Live-cell imaging of Gfp-Rpb4 in the wild-type and *rtr1Δ* mutant strains. **D**) Live-cell of Rpb1-Gfp localization in the wild-type and *rtr1Δ* mutant strains.

Although these data suggested that the association between Rpb1 and Rpb3 (and likely Rpb2) was not altered, the relative depletion of heterodimer Rpb4/7 in relation to the remaining RNA pol II has been observed in purification experiments using Rtr1-TAP as bait [34]. Furthermore, a strong genetic interaction between *RTR1* and *RPB4* has been reported [27]. Based on these data and ours above, we explored if lack of Rtr1 would affect the association of Rpb4 with the rest of RNA pol II. Accordingly, we analysed by western blot the Rpb4/Rpb1 ratio in the above-indicated immunoprecipitation assays (Figure 1A and S1A). Notably, a significant decrease in the association between Rpb4 and Rpb1 was observed. Then we extended this analysis to other RNA pol II small subunits like Rpb7, which forms a heterodimer with Rpb4, and also Rpb5 (Figures 1A and S1A). As these results demonstrate, and as expected, the association between Rpb7 and Rpb1 also decreased, which suggests the need of Rtr1 for the correct association of dimer Rpb4/7 with the rest of the RNA pol II. Interestingly, Rpb5/Rpb1 also dropped in the *rtr1Δ* mutant cells.

According to the proposed model for the biogenesis of eukaryotic multisubunit RNA polymerases [14], we speculated that the defect in the Rpb5 association could be a result of deficiencies in the formation of the subassembly complex Rpb1 (composed of Rpb1, Rpb5, Rpb6 and Rpb8), which must interact with subassembly complexes Rpb2 and Rpb3 to form the complete enzyme prior to its nuclear import [14].

To test this hypothesis, we used an *rtr1Δ* mutant and its isogenic wild-type strain, which both express a functional Rpb8-TAP tagged protein, and performed immunoprecipitation with the Rpb3 subunit of RNA pol II prior to the western blot to analyse the subunits’ RNA pol II composition (Rpb5, Rpb6 and Rpb8) with specific antibodies against, or with the anti-TAP antibody for Rpb8-TAP. We also analysed Rpb4, Rpb7, Rpb1 and Rpb3. As shown in Figures 1B and S1B, in this new genetic background the results shown above for the Rpb3/Rpb1, Rpb4/Rpb1 and Rpb7/Rpb1 ratios were confirmed. Notably the western blot analysis showed lower levels of subunits Rpb5, Rpb6, and Rpb8 *vs*. Rpb1 in the mutant strain compared to its wild-type counterpart (Figure 1B and S1B). As a control, the Rpb8-TAP pull down from the *rtr1Δ* mutant and its isogenic wild-type strain corroborated the decrease in Rpb4 *vs*. Rpb1 in the mutant strain (Figure S1C).

The deletion of *RTR1* has been described to cause the partial cytoplasmic mislocalization of Rpb1 [26]. Furthermore, silencing of RPAP2, the human ortholog of Rtr1, also leads to Rpb1 accumulation in the cytoplasm of human cells [38]. Accordingly, we analysed if *RTR1* deletion would also provoke the cytoplasmic accumulation of Rpb4. To do so, we employed the *rtr1Δ* mutant and wild-type strains that express Rpb4-GFP, and analysed Rpb4 by fluorescent microscopy. As shown in Figure 1C, Rpb4 clearly accumulated in both the cytoplasm and nucleus of *rtr1Δ* mutant cells, while only nuclear localization was observed in the wild-type cells. As a control, we analysed Rpb1 localization in the *rtr1Δ* mutant and wild-type strains that express Rpb1-GFP, and corroborated the Rpb1 mislocalization previously described under Rtr1 inactivation [26] (Figure 1D).

Taken together, our data suggest a role for Rtr1 in RNA pol II assembly, likely in the cytoplasm, probably by acting as an assembly factor needed for both the correct formation of subassembly complex 1 and the association of heterodimer Rpb4/7 with the rest of the enzyme.

### The assembly of RNA pol I and III does not alter under *RTR1* deletion

Our above results demonstrated that the assembly of Rpb5, Rpb6 and Rpb8 with RNA pol II was impaired in the *rtr1Δ* mutant cells. As these proteins are three of the five common subunits shared by the three RNA polymerases in eukaryotes (reviewed in [59]), we speculated that Rtr1 could also participate in the assembly of RNA pols I and III. To explore this in detail, we used the *rtr1Δ* mutant and wild-type strains expressing Rpa190::HA or Rpc160::Myc, the tagged forms of the largest subunits of RNA pol I and RNA pol III, respectively. By using specific antibodies against the tags, we performed protein immunoprecipitation and analysed Rpb5 co-immunoprecipitation. As seen in Figure 2, no differences in the Rpb5/Rpa190 or Rpb5/Rpc160 ratios were observed in the *rtr1Δ* mutant cells in relation to the wild-type counterparts, which suggests that assembly of RNA pols I and III was not impaired under *RTR1* deletion.

**Figure 2:**
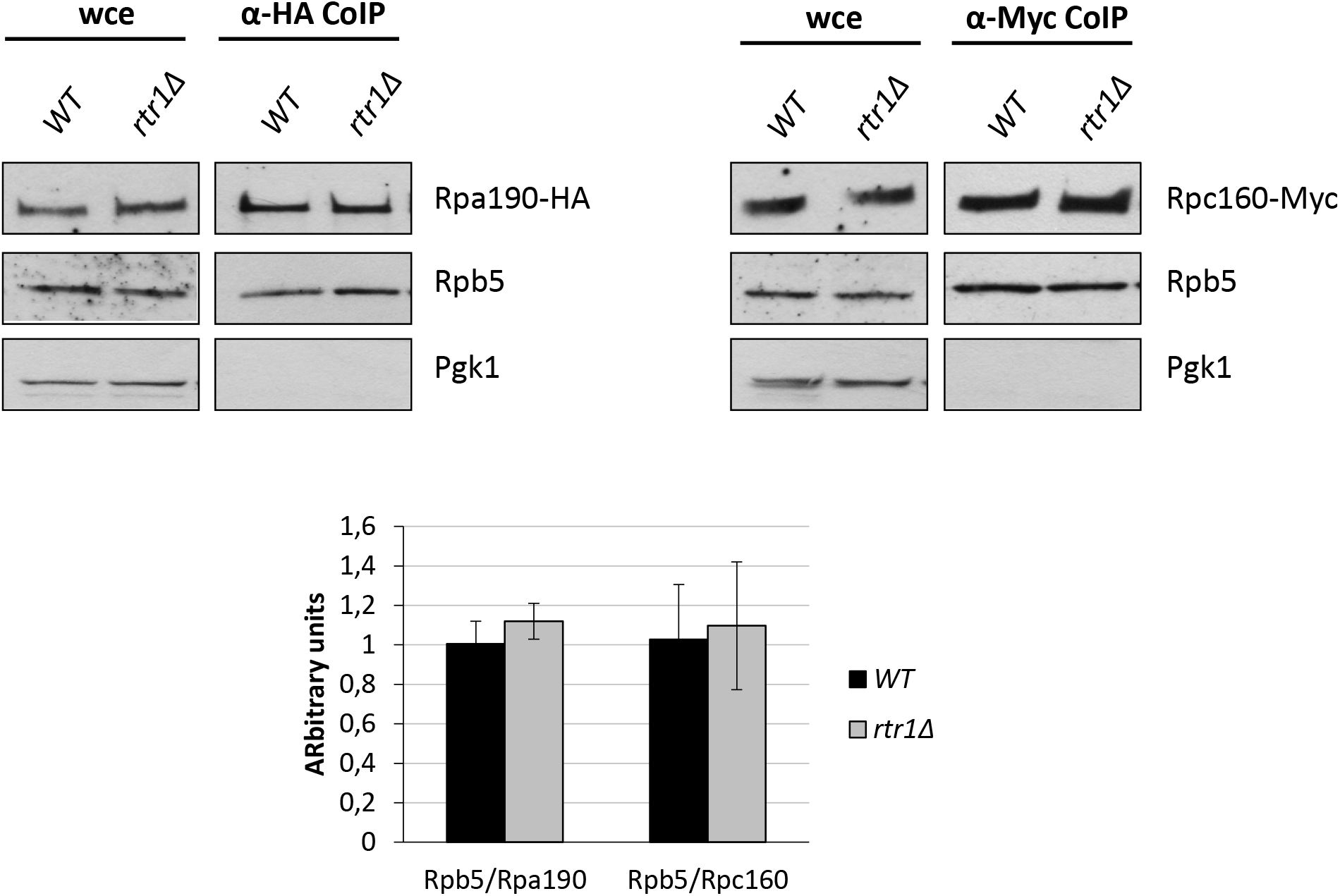
*RTR1* deletion does not affect the assembly of RNA pol I and III. Left panel: western blot of the whole cell crude extracts and RNA pol I immunoprecipitation (using an HA antibody that recognized an HA tagged form of Rpa190) from the wildtype and *rtr1Δ* mutant strains. Right panel: western blot from the whole cell crude extracts and RNA pol III immunoprecipitation (using a cMyc specific antibody that recognized the C160-myc protein) from the *rtr1Δ* mutant and its isogenic wild-type strains. Cells were grown in YPD medium at 30°C. Lower panel, the quantification of western blot showing the Rpb5/Rpa190 and Rpb5/Rpc160 ratios in the *rtr1Δ* mutant and its wild-type isogenic strain. Median and SD of at least two independent biological replicates.

All these data point out that Rtr1 specifically acts on RNA pol II assembly, but not on the assembly of RNA pols I and III.

### Rtr1 and the prefoldin-like Bud27 may cooperate for RNA pol II assembly

Our data above suggest the functional relation between Rtr1 and Rpb5 in RNA pol II assembly by showing that *rtr1Δ* mutant cells reduced the association of Rpb5 with RNA pol II. Notably, prefoldin-like Bud27, and its human ortholog URI, were associated with Rpb5 [22,60,61]. Furthermore, Bud27 mediates the assembly of the three RNA pols in an Rpb5-dependent manner [22,23]. URI also participates in RNA pol II cytoplasmic assembly [15,16,22,24] and has been described as a part of the HSP90/R2TP-prefoldin-like complex found in the subcomplexes with RPB5 [15,16,24].

We wondered if Rtr1 could act in concert with prefoldin-like Bud27. Accordingly, we firstly analysed the genetic interactions between *RTR1* and *BUD27*. As shown in Figure 3A, *BUD27* overexpression strongly aggravated the formamide sensitivity phenotype of the *rtr1Δ* mutant cells. Furthermore, the *rtr1Δbud27Δ* double mutant exhibited a higher temperature sensitivity phenotype than the corresponding single mutants (Figure 3B). To investigate if the genetic interactions between *RTR1* and *BUD27* could also reflect a functional interaction, we explored whether lack of Rtr1 would affect the physical association between Bud27 and RNA pol II. To do so, we performed Rpb3 immunoprecipitation in the *rtr1Δ* mutant and the wild-type strains expressing a functional Bud27 TAP-tagged version of this protein, and analysed Bud27 coimmunoprecipitation. As seen in Figure 3C, no significant differences between Bud27 and the Rpb1 ratio were observed in the cells lacking Rtr1, which indicates that Rtr1 did not seem to affect the association between Bud27 and RNA pol II. As a control, we corroborated the significant drop in the Rpb5/Rpb1 ratio (Figure 3C). In an effort to decipher the functional relation between Rtr1 and Bud27, we also performed Rpb3 immunoprecipitation in the *bud27Δ* mutant and wild-type strains, both expressing functional Rtr1 TAP-tagged. Accordingly to the results above, *BUD27* deletion did not affect the association of Rtr1 with Rpb1, while a significant drop in the Rpb5/Rpb1 ratio was observed, which was expected given the role of Bud27 in RNA polymerase assembly [22] (Figure 3D).

**Figure 3:**
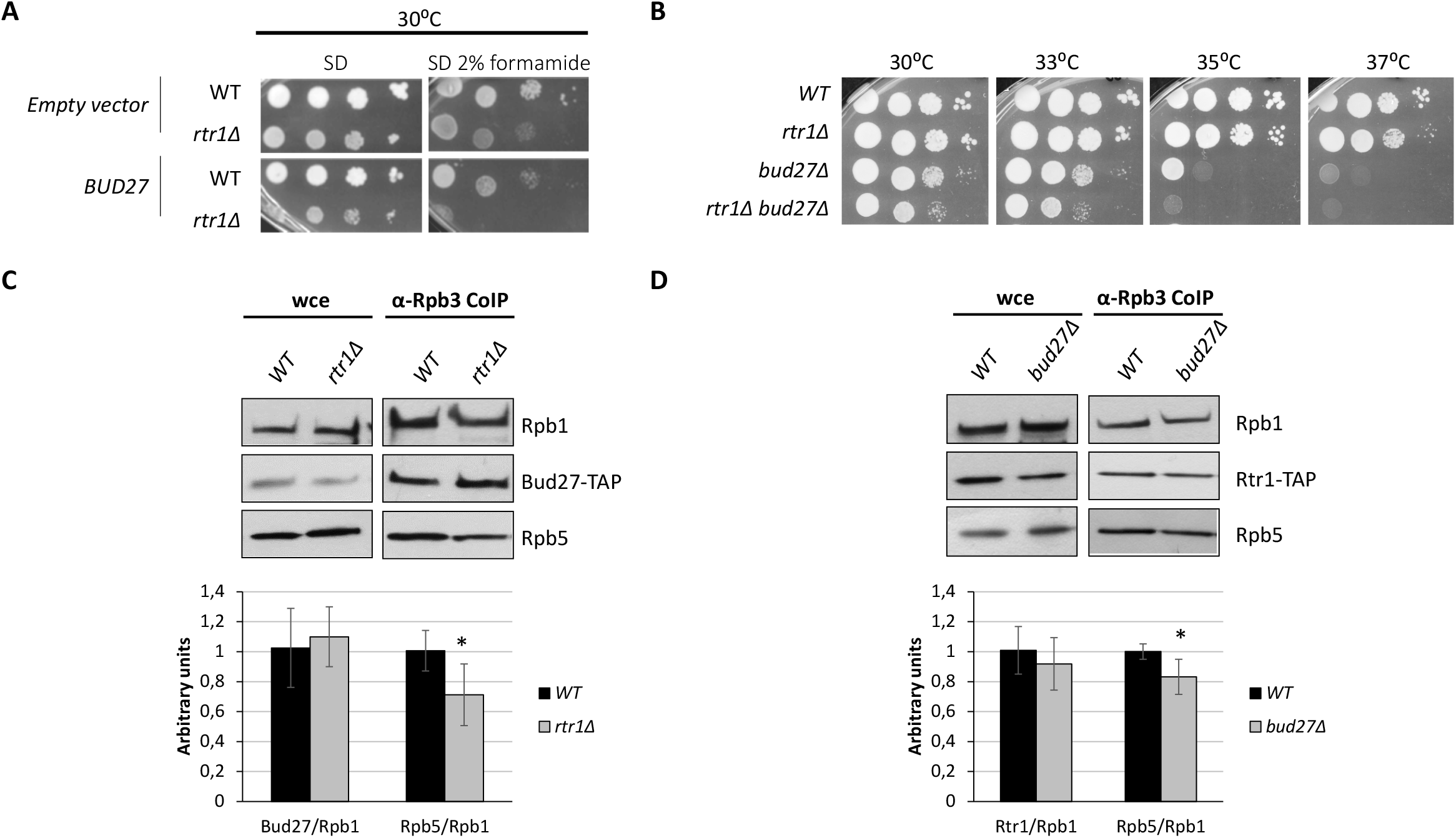
Analysis of the Rtr1 and Bud27 functional relation. **A**) Analysis of *BUD27* overexpression in the wild-type and *rtr1Δ* mutant cells growing in SD medium or SD supplemented with 2% formamide at 30°C. **B**) Genetic interaction of *rtr1Δ* and *bud27Δ* mutants. Growth of single and double mutants in YPD medium at different temperatures. **C**) RNA pol II immunoprecipitation using an Rpb3 antibody from the wild-type and *rtr1Δ* mutant cells containing a *bud27*::TAP allele grown in YPD medium at 30°C. Lower panel: the Bud27/Rpb1 and Rpb5/Rpb1 ratios from the quantifications of western blots. Median and SD of at least two independent biological replicates. The Rpb1 anti-CTD antibody was used. **D**) RNA pol II immunoprecipitation using an Rpb3 antibody from the wild-type and *bud27Δ* mutant cells containing a functional Rtr1-TAP tag. Lower panel: the Rtr1/Rpb1 and Rpb5/Rpb1 ratios from the quantifications of western blots. Median and SD of at least two independent biological replicates. *p<0.05 (t-test).

These data indicate a functional relation between Bud27 and Rtr1 that seems to be independent of their association with RNA pol II.

### Lack of Rtr1 affects the chromatin-associated RNA Pol II composition

Our data revealed that lack of Rtr1 affected the RNA pol II assembly. Lack of Rtr1 also affects RNA pol II import to the nucleus [26,38]. To explore whether this assembly defect could impact the RNA pol II complexes associated with chromatin, likely engaged in transcription, we analysed the subunits’ composition of nuclear RNA pol II associated with chromatin. For this purpose, we took advantage of the yChEFs procedure [58,62] to isolate chromatin-enriched fractions with their associated proteins in both *rtr1Δ* cells and the wild-type counterpart, both expressing a functional TAP-tagged Rpb8 (Figure 4 and S2). The chromatin fractions obtained by yChEFs were used to analyse RNA pol II composition by western blot with specific antibodies against some RNA pol II subunits, or against anti-TAP for Rpb8-TAP. By employing specific antibodies against 3-phosphoglycerate kinase (Pgk1) as a control of cytoplasmic protein, and against histone-3, as a control of chromatin-associated proteins, we observed that chromatin was successfully isolated and similar chromatin levels were purified and analysed in the wild-type and mutant cells. Notably, and unlike that observed when purifying the global RNA pol II, the levels of Rpb1, Rpb3, Rpb5, Rpb6, Rpb7 and Rpb8 associated with chromatin were similar in both the wild-type and *rtr1Δ* mutant cells. These results suggested that the non-assembled and nonfunctional RNA pol II subcomplexes lacking some of these subunits did not bind chromatin and likely did not shuttle to the nucleus and continued to accumulate in the cytoplasm.

**Figure 4:**
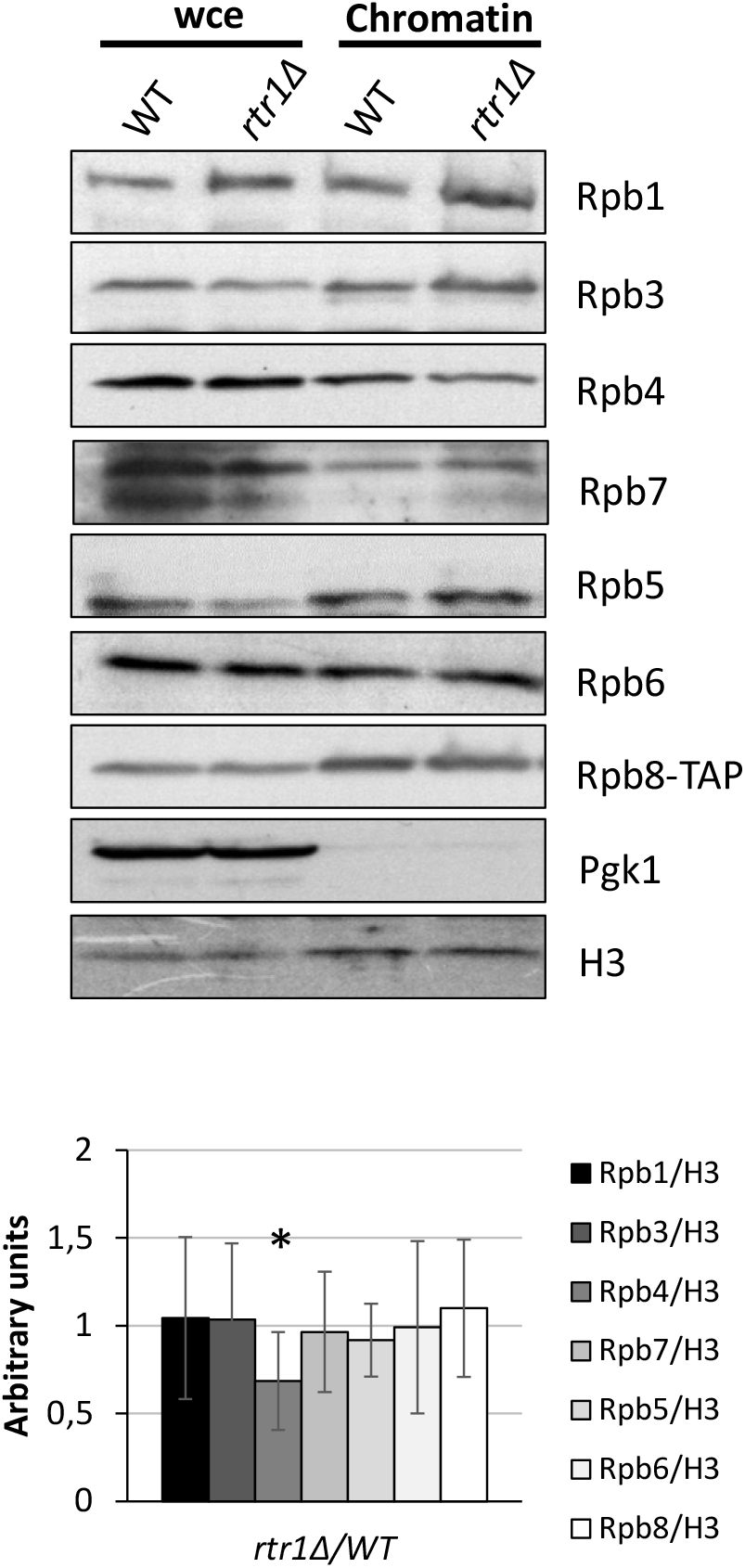
Analysis of chromatin-associated RNA pol II. Whole-cell extract and chromatin-associated proteins isolated by the yChEFs procedure [58,62] from the wild-type and *rtr1Δ* strains containing an Rpb8-TAP tag, grown in YPD medium at 30°C, were analysed by western blot with specific antibodies against Rpb1 (y-80), Rpb3, Rpb4, Rpb7, Rpb5, Rpb6, and TAP tagged Rpb8. H3 histone was used as a positive control of the chromatin-associated proteins, and Pgk1 was the negative control of cytoplasmic contamination. The quantification for each RNA pol II subunit *vs*. histone H3 between the *rtr1Δ* mutant and wild-type strains. Median and SD of at least three independent biological replicates. *p<0.05 (t-test).

However, the amount of Rpb4 associated with chromatin significantly diminished in the *rtr1Δ* strain in relation to the wild-type counterpart (Figure 4), as did the Rpb4/Rpb1 ratio (Figure S2). These results suggest that two RNA pol II populations bound chromatin when Rtr1 was lacking: complete RNA pol II and RNA pol II lacking Rpb4. Notably, we observed *RTR1* as a multicopy suppressor of the temperature sensitivity phenotype of *RPB1* mutant *rpo21-4*, which provokes the dissociation of dimer Rpb4/7 from the rest of the enzyme [63] (Figure 5A), corroborating the abovementioned conclusion. In addition, *RTR1* deletion is lethal when combined with both *rpo21-4* and the *rpb1-84* foot mutation, with similar consequences on RNA pol II assembly [63]. We also observed genetic interactions between the *rtr1Δ* mutant and the *rpb6Q100R* mutant, which also caused loss of dimer Rpb4/7 from the rest of RNA pol II [64] (Figure 5B). Finally, in agreement with the effect provoked by lack of Rtr1 on RNA pol II assembly, *rtr1Δ* showed a strong genetic interaction with not only the *rpb7ΔC3* mutant of the Rpb7 subunit (Figure 5C), but also with the *rpb4Δ* mutant, as previously reported [27] (Figure 5C).

**Figure 5:**
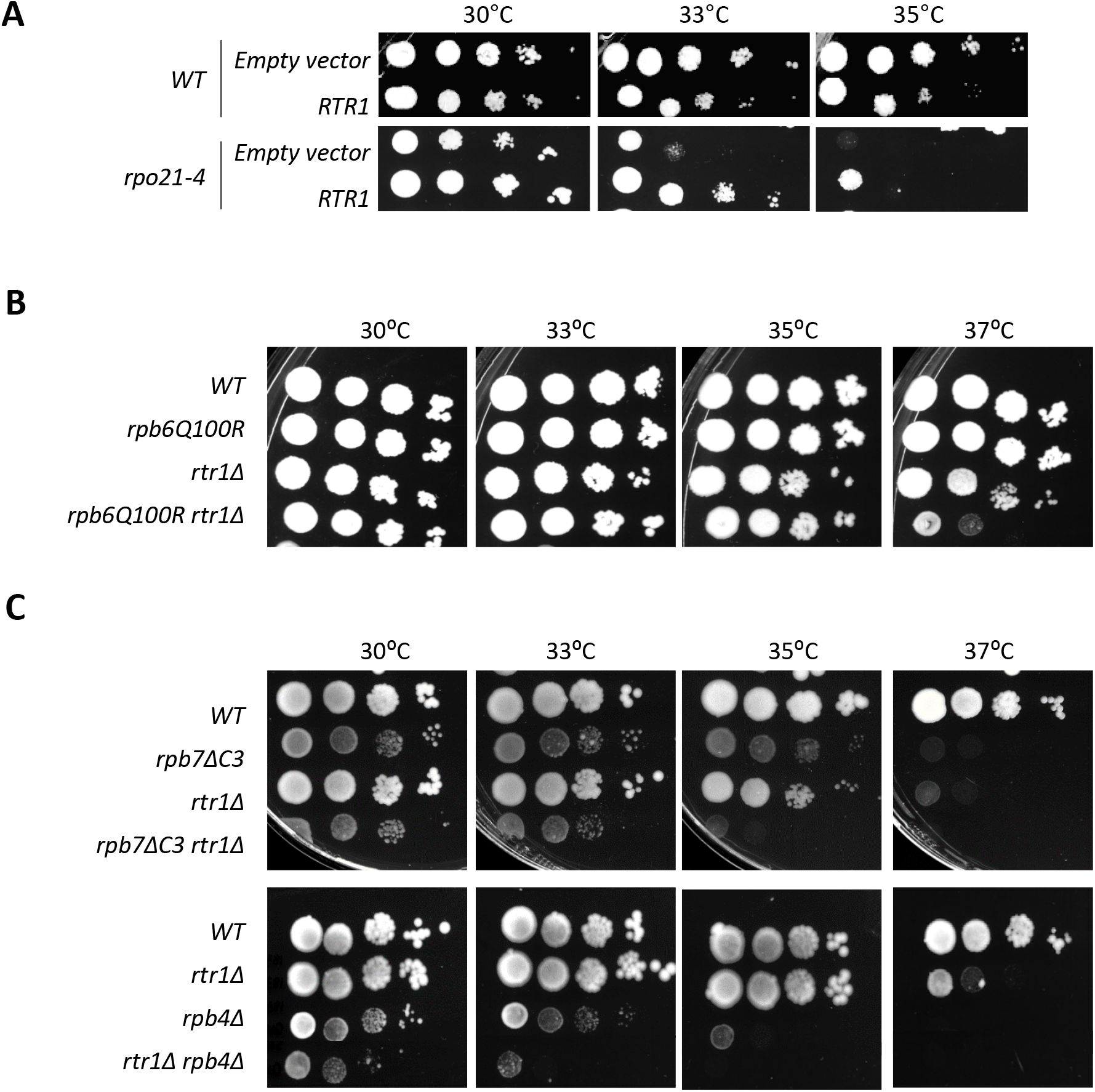
Genetic interactions between *RTR1* and *RPB1, RPB6, RPB4* and *RPB7*. **A**) *RTR1* overexpression in the *rpo21-4* and *rpb1-84* foot mutants (*RPB1* mutations) grown in SD at different temperatures. **B**) *rtr1Δ* and *rpb6Q100R* single and double mutants grown at different temperatures in YPD medium. **C**) Growth of the *rtr1Δ, rpb7ΔC3* and *rpb4Δ* single and double mutants in YPD medium at different temperatures.

Based on these data, we point out that *RTR1* deletion provokes a defect in RNA pol II assembly and leads to the cytoplasmic accumulation of RNA pol II subcomplexes, and probably of free RNA pol II subunits. Concomitantly, the nuclear import of complete RNA pol II and RNA pol II lacking Rpb4, and likely dimer Rpb4/7, would occur, which would associate with chromatin to be engaged in transcription. In any case, we cannot rule out two scenarios: the Rpb4 association with RNA pol II could also occur in the nucleus and be affected by lack of Rtr1; an impact on Rpb4 dissociation during transcription in the *rtr1Δ* mutant cells in line with previous data [34].

### Lack of Rtr1 does not account for a global effect on Rpb4 dissociation from RNA pol II during transcription elongation

Previous work by quantitative proteomic analyses states that Rpb4 and Rpb7 dissociate from the rest of RNA pol II during transcription elongation [34], likely with the participation of Rtr1, whose association with the enzyme requires Ser2 CTD phosphorylation [41].

To investigate whether the decrease in Rpb4 associated with RNA pol II that we observed when Rtr1 was lacking could be the consequence of a defect in Rpb4 dissociation during transcription, we explored by ChIP Rpb3 and Rpb4 occupancy along the entire length of different transcriptional units in the wild-type and *rtr1Δ* mutant cells. As shown in Figure 6, Rpb3 occupancy increased from the promoter region to halfway or the beginning of *PMA1* and *PYK1* ORFs, respectively, and decreased through the 3’ regions in the wild-type and *rtr1Δ* mutant cells despite lack of Rtr1 diminishing Rpb3 occupancy for the entire transcriptional units in relation to the wild-type cells. These results fall in line with those previously reported [29]. Notably, Rpb4 occupancy showed a similar profile to Rpb3 occupancy in the wild-type and *rtr1Δ* mutant cells, and peaked at the beginning of genes *PMA1* and *PYK1*, before decreasing through the 3’ regions (Figure 6). However, Rpb4 occupancy drastically decreased for each analysed region when Rtr1 was lacking in relation to the wild-type cells (Figure 6). The dissociation of Rpb4 from the rest of RNA pol II during transcription was represented by the ratio between Rpb4 and Rpb3 occupancies along the entire length of the transcriptional units. As shown in Figure 6, the Rpb4/Rpb3 ratios for *PMA1* and *PYK1* lowered from the beginning to the 3’ region of these genes in the wild-type cells, which supports Rpb4 dissociation occurring [34,65,66]. Strikingly, no major differences in the Rpb4/Rpb3 profile along the *PMA1* and *PYK1* transcription units were observed in the *rtr1Δ* mutant cells in relation to the wild-type strain, which suggests that lack of Rtr1 did not significantly impact the global Rpb4 dissociation. However, as expected from data above, the Rpb4/Rpb3 ratios lowered for each analysed region, which corroborates that Rpb4 deficiency *vs*. the rest of RNA pol II observed in chromatin fractions was the consequence of a defect in Rpb4 association with the rest of the enzyme, and not of Rpb4 dissociation during transcription. Our results do not exclude this association from occurring, at least partly, in the nucleus. Similar results, albeit with differences in the Rpb3 and Rpb4 occupancy profiles between the wild-type and *rtr1Δ* mutant cells were also observed for the *URA2* transcription unit (Figure 6). Finally, we did not observe any decrease in Rpb4/Rpb3 ratio for the *MTG1* gene (Figure S3), which could fall in line with data that propose the influence of Rtr1 and other elongation factors on Rpb4 dissociation during transcription for a limited number of genes [34].

**Figure 6:**
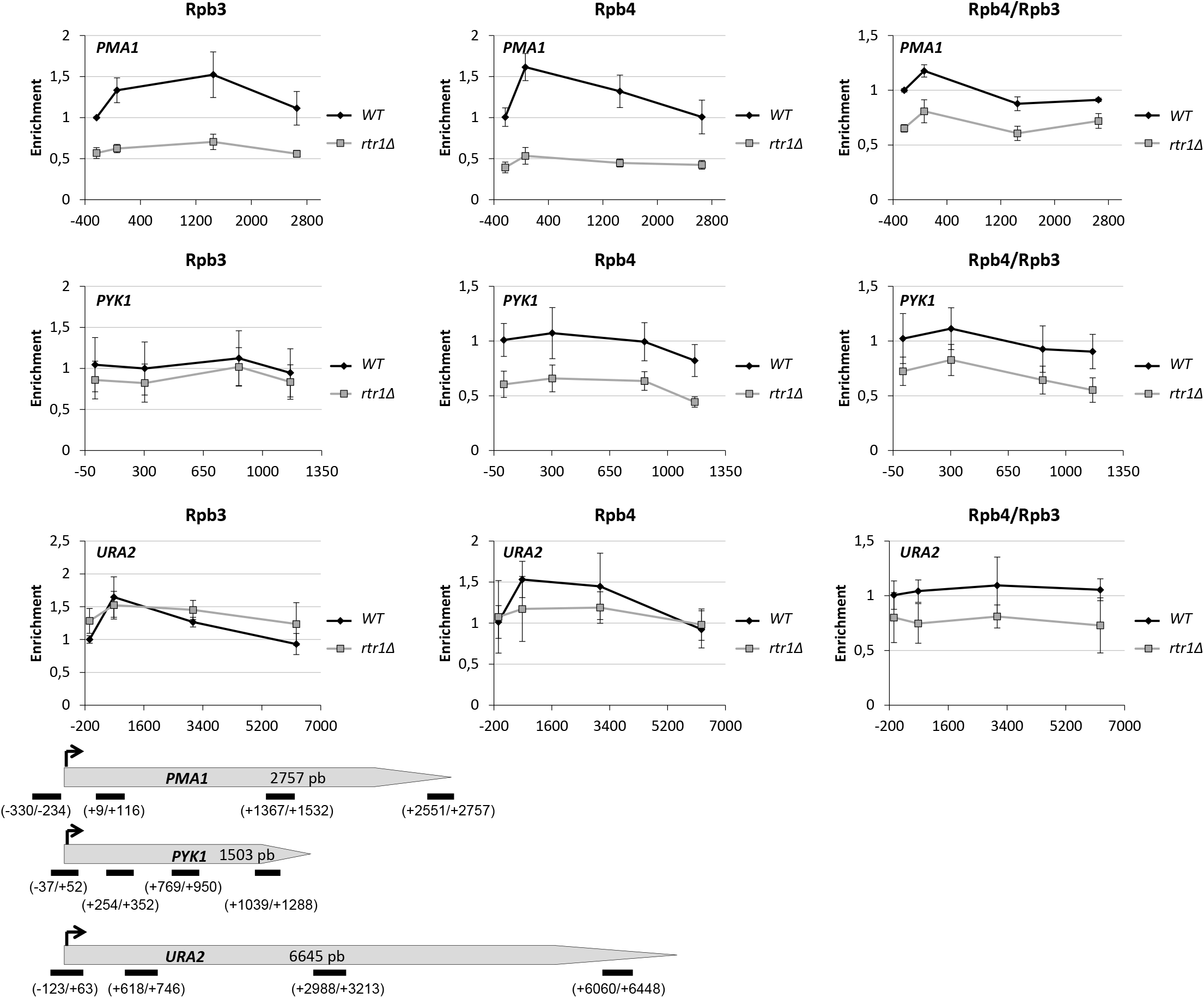
The *rtr1Δ* mutation affects gene occupancy by RNA pol II, but not global Rpb4 dissociation. Chromatin immunoprecipitation (ChIP) analysis for different genes in the wild-type and *rtr1Δ* cells, performed with anti-Rpb3 (left panels) and anti-Rpb4 (middle panels) antibodies, against the Rpb3 and Rpb4 RNA pol II subunits. Right panels: Rpb4/Rpb3 ratios for the Rpb4 dissociation analysis, from the left and middle panel’s results. Lower panel: the transcription units used in this work indicating the location of the analysed PCR amplicons. The values found for the immunoprecipitated PCR products were compared to those of the total input, and the ratio of each PCR product of transcribed genes to a non-transcribed region of chromosome V was calculated.

### Incorrect RNA pol II assembly in the *rtr1Δ* mutant affects mRNA stability

It has been demonstrated that Rpb4 imprints mRNA, in the context of transcribing RNA pol II, and regulates mRNA decay in an Rpb7-dependent manner. Consequently, the decrease in Rpb4-mRNA imprinting increases mRNA stability [48,53,55,56,67,68,69].

Accordingly, we speculated that the defect in RNA pol II assembly provoked by lack of Rtr1 could impact Rpb4-mRNA imprinting and, consequently, mRNA stability. To investigate this possibility, we firstly analysed whether the association of Rpb4 with mRNAs could alter in *rtr1Δ* mutant cells. To do so, we crosslinked the proteins bound to mRNA by using UV irradiation at 254 nm, as previously described [48], from *rtr1Δ* mutant and wild-type cells. Having isolated total poly(A)-containing mRNA, the association of Rpb4 with mRNAs was analysed by western blot with specific antibodies. As shown in Figure 7A, *RTR1* deletion reduced the amount of Rpb4 associated with mRNA in relation to its wild-type counterpart, which indicates that Rpb4-mRNA imprinting is influenced by Rtr1. As a control, as shown by analysing the presence of the Pgk1 protein in the samples with a specific antibody, no major cytoplasmic contamination was detected.

**Figure 7:**
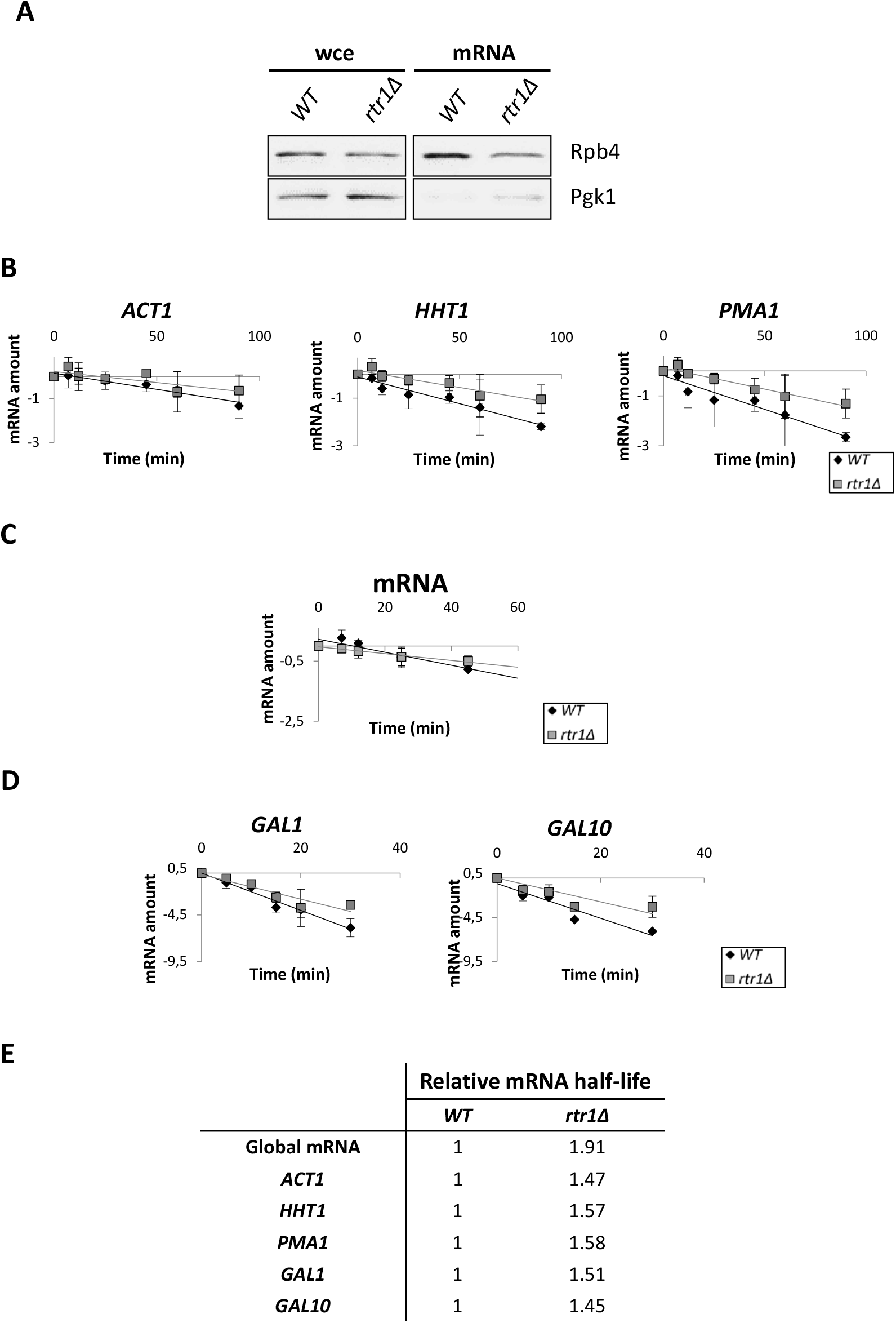
*rtr1Δ* decreases Rpb4-mRNA imprinting and increases mRNA stability. **A**) Western-blot of Rpb4 in both whole-cell free extracts and oligo-dT purified mRNAs after exposure to 1,200 mJ/cm^2^ of 254 nm UV in the wild-type and *rtr1Δ* mutant. Anti-Pgk1 antibody was used as a negative control. **B**) mRNA levels, measured by RT-qPCR, for genes *ACT1, HHT1* and *PMA1* after thiolutin addition to block transcription in the wild-type and *rtr1Δ* mutant. **C**) Global mRNA levels measured at different times after thiolutin addition to block transcription in the wild-type and *rtr1Δ* mutant using an oligodT probe by a dot-blot assay. **D**) *GAL1* and *GAL10* mRNA amounts measured by RT-qPCR, in the wild-type and *rtr1Δ* mutant at different times after blocking transcription with glucose. Time 0 corresponded to the cells grown in the presence of galactose as the carbon source. In B, C and D, the drop in the mRNA levels after shutoff at different times is represented on a natural logarithmic scale. In B and D, rRNA 18S was used as a normalizer. E) The relative mRNA half-lives calculated from the experiments represented in B, C and D. All the experiments corresponded to at least three independent biological replicates.

To explore if the reduction in Rpb4-mRNA imprinting affected mRNA stability, we proceeded as in [48] and we treated the cultures of the wild-type and *rtr1Δ* cells with 5 μg/ml thiolutin to block transcription. Total RNA was extracted at different times after thiolutin addition. The RT-qPCR analysis of the mRNA levels for genes *ACT1, HHT1* and *PMA1* demonstrated an increase in the mRNA half-live for all the tested genes in the *rtr1Δ* mutant in relation to the wild-type strain (Figure 7B and 7E). The global mRNA decay analysis by dot-blot using a fluorescent oligo-dT probe corroborated these results by revealing a 1.91-fold global half-life increase in the mutant *versus* the wild-type strain (Figure 7C and E). Finally, these data were also confirmed by a different transcription shutoff, by analysing *GAL1* and *GAL10* genes mRNA decay in the cells growing in SD medium containing galactose as a carbon source, which were then shifted to glucose as the sole carbon source to stop transcription (Figure 7D and E).

All these data indicate that the alteration of RNA pol II assembly provoked by *RTR1* deletion would affect the amount of Rpb4-containing RNA pol II associated with chromatin, and would consequently decrease Rpb4-mRNA imprinting, increasing mRNA stability. Indeed similar results have been previously demonstrated for other *rpb1* and *rpb6* mutants where defects in RNA pol II integrity alters the correct dimer Rpb4/7 association [48,54,55,63].

### *RPB5* overexpression restores the assembly of RNA pol II in *rtr1Δ* cells, and partially overcomes mRNA stability

In order to demonstrate that the mRNA stability defects of *rtr1Δ* cells are directly associated with incorrect RNA pol II assembly, we planned to seek a situation in which RNA pol II assembly defects are suppressed. Accordingly, we firstly overexpressed the genes for some RNA pol II subunits altered by *RTR1* deletion: *RPB5, RPB4, RPB4 + RPB7* and *RPB6*. As shown in Figure 8A, only *RPB5* overexpression overcame the sensitivity to 2% formamide phenotype of the *rtr1Δ* mutant, as previously described [27]. However, and surprisingly, neither *RPB4* nor *RPB4 + RPB7* overexpression suppressed the above-mentioned phenotype, nor did *RPB6* overexpression, the gene coding for the Rpb6 subunit which connects dimer Rpb4/7 with the core of RNA pol II [22,63,70].

**Figure 8:**
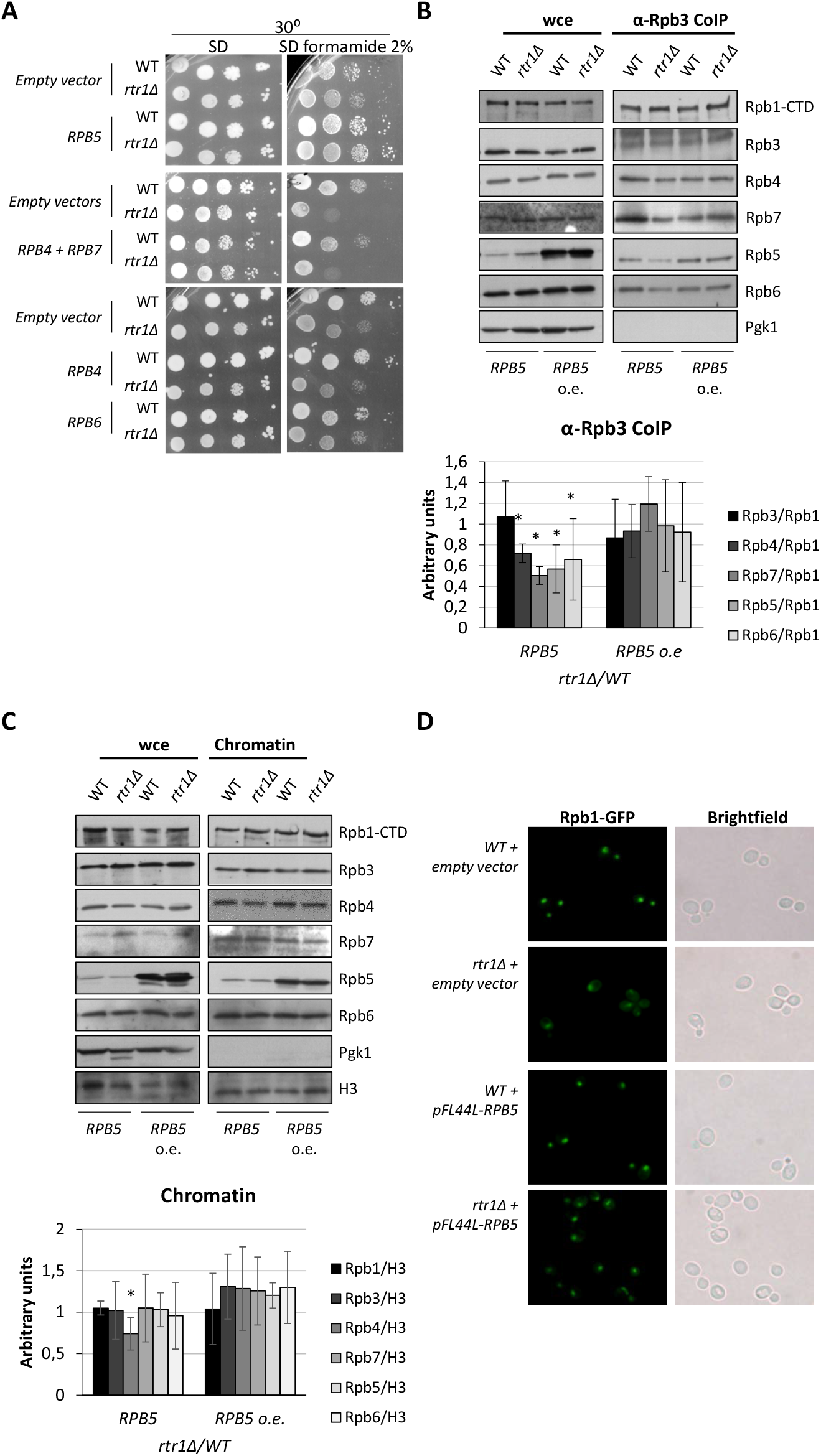
*RPB5* overexpression overcomes the RNA pol II assembly defect in the *rtr1Δ* mutant. **A**) *RPB4, RPB5, RPB6* and *RPB4/7*overexpression in the wild-type and *rtr1Δ* mutant cells grown in SD and SD supplemented with 2% formamide at 30°C. **B**) Rpb3 immunoprecipitation in the *rpb5Δ* and *rpb5Δ rtr1Δ* cells both expressing the *RPB5* gene from a centromeric (*RPB5*) or a high copy number (*RPB5* o.e.) plasmid, and grown in YPD medium at 30°C. The western blots of Rpb1, Rpb3, Rpb4, Rpb7, Rpb5, and Rpb6 from the whole cell crude extracts and immunoprecipitated samples are shown (upper panel). Pgk1 was used as a negative control of the RNA pol II immunoprecipitated samples. Lower panel: the quantification of the western blot showing the *rtr1Δ* mutant/wild-type strains ratio for each subunit *vs*. Rpb1. *p<0.05 (t-test). Median and SD of at least three independent biological replicates. **C**) The wholecell extract and chromatin enriched fractions obtained by the yChEFs procedure [58,62] from the same strains used in B. Cells were grown in YPD medium at 30°C and proteins were analysed by western blot with specific antibodies against Rpb1, Rpb3, Rpb4, Rpb7, Rpb5 and Rpb6. H3 histone was used as a positive control of the chromatin-associated protein and Pgk1 employed as a negative control of cytoplasmic contamination. Lower panel: the quantification for each RNA pol II subunit *vs*. histone H3 between the *rtr1Δ* mutant and wild-type strains. Median and SD of at least three independent biological replicates. *p<0.05 (t-test). **D**) Rpb1-GFP localization in the *rtr1Δ* mutant and its wild-type counterpart after *RPB5* overexpression using a high copy number plasmid.

Then we wondered whether *RPB5* overexpression could restore the RNA pol II assembly defects observed in the *rtr1Δ* cells. To do so, we immunoprecipitated Rpb3 from the *rtr1Δ rpb5Δ* double mutant strain and its *rpb5Δ* counterpart, both complemented with a multicopy plasmid overexpressing *RPB5* or a centromeric plasmid expressing *RPB5* from its own promoter (note that *RPB5* gene is essential). After immunoprecipitating RNA pol II, we analysed by western blot the coimmunoprecipitation of other RNA pol II subunits. Figure 8B shows how the *rtr1Δ rpb5Δ* mutant cells overexpressing *RPB5* overcame the assembly defect of subunits Rpb4, Rpb5, Rpb6 and Rpb7 of the *rtr1Δ rpb5Δ* mutant cells expressing *RPB5* from a centromeric plasmid. Furthermore, and notably, the correction of RNA pol II assembly defect by *RPB5* overexpression led to wild-type chromatin-associated RNA pol II levels in the *rtr1Δ* mutant cells, overcoming the alteration in Rpb4 association (Figure 8C, compared to Figures 4 and S2). In addition, *RPB5* overexpression partially suppressed the Rpb1 mislocalization observed in the *rtr1Δ* mutant (Figure 8D).

As *RPB5* overexpression restored RNA pol II assembly and the decrease in Rpb4 binding to chromatin-associated RNA pol II in the *rtr1Δ* cells, we wondered if this suppression would suffice to also restore mRNA stability. To answer this question, we used a multicopy plasmid to overexpress *RPB5* in not only the *rtr1Δ* mutant cells, but also in its isogenic wild-type strain. As a control, the same strains were transformed with an empty vector. The transformed cells were grown in SD medium containing galactose as the carbon source to induce *GAL1* gene expression. Then *GAL1* gene repression was monitored by shifting cells from galactose-to glucose-containing medium to stop *GAL1* expression, and mRNA stability was analysed at different times. As shown in Figure 9, *RPB5* overexpression partially suppressed the mRNA stability alteration that occurred to the *rtr1Δ* mutant cells (about 20 %).

**Figure 9:**
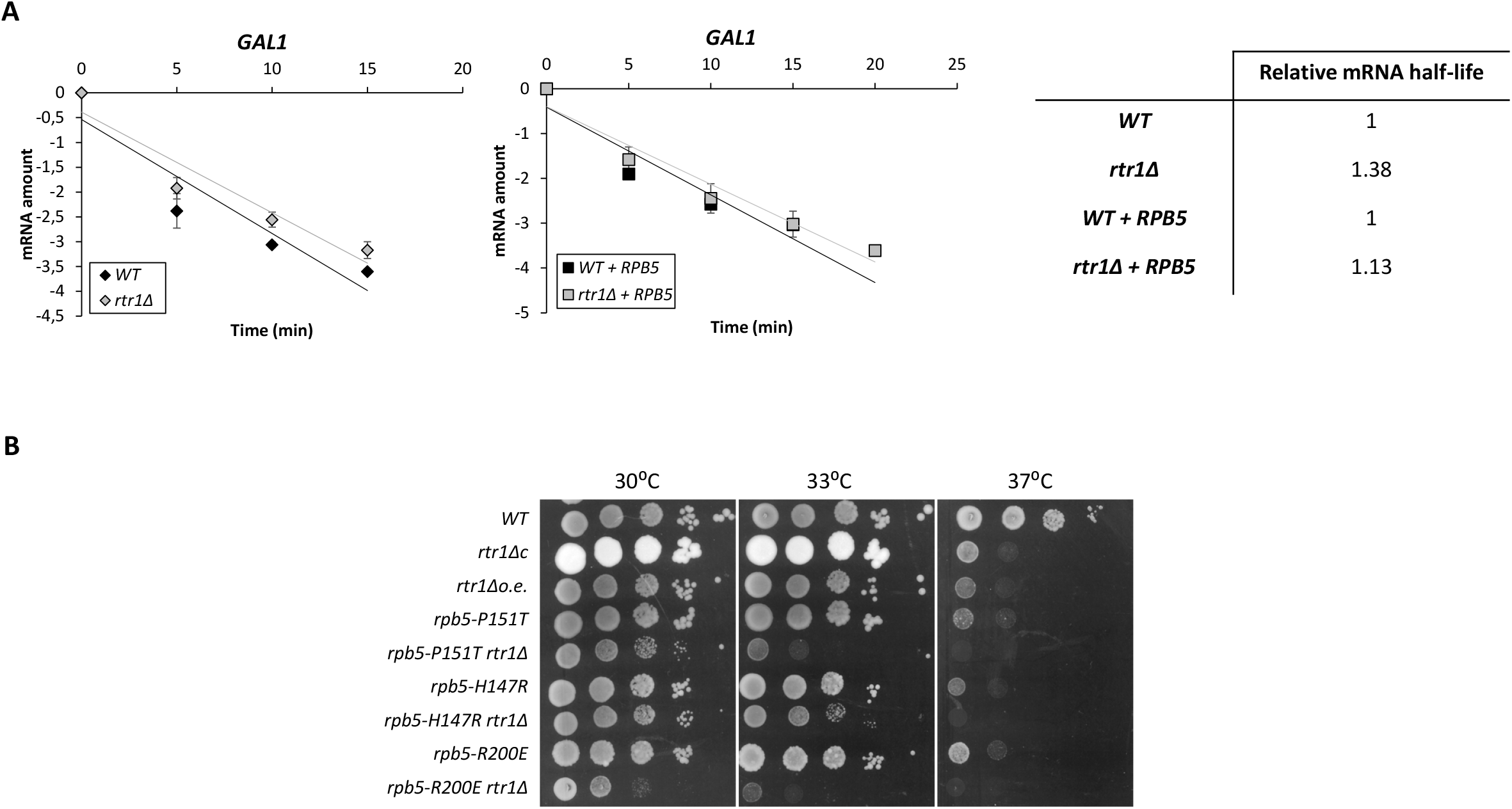
*RPB5* overexpression overcomes partially the mRNA stabilisation of the *rtr1Δ* mutant. **A**) *GAL1* mRNA amount in the wild-type and *rtr1Δ* mutant transformed with a high copy number plasmid containing *RPB5* and the same empty vector used as a control. *GAL1* mRNA amount was measured by RT-qPCT at different times after blocking transcription with glucose using rRNA 18S as a normalizer. Time 0 corresponds to the cells grown in the presence of galactose as the carbon source. The drop in the mRNA levels after shutoff at different times is represented on a natural logarithmic scale. Right panel: the relative mRNA half-lives calculated from the RT-qPCR experiments. Median and SD of at least three independent biological replicates. **B**) Genetic interactions between the *rtr1Δ* mutant and different *rpb5* mutants. Single and double mutants were grown in YPD at different temperatures. Wild-type (WT) corresponds to the YFN2 strain bearing the *rpb5Δ::ura3::LEU2* allele and complemented with multicopy plasmid *pFL44L-RPB5 (2 μm URA3 RPB5*). The rest of the strains are YFN2 derivative. The single *rtr1Δ* mutant strains are also YFN2 derivative expressing the *RPB5* gene from a centromeric (*rtr1Δc*) or a high copy number (*rtr1Δ*o.e.) plasmids.

Finally, in line with the functional interaction between Rpb5 and Rtr1, we also observed genetic interactions between genes *RPB5* and *RTR1*, as shown by the phenotype analysis of the double *rpb5* mutant alleles in combination with the *rtr1Δ* mutation in relation to the single mutants and wild-type strains (Figure 9B).

The above data suggest that overcoming the defect in Rpb4 binding to the chromatin-associated RNA pol II caused by lack of Rtr1 also partially suppressed the mRNA stability defect. However, *RPB4/7* overexpression overcame neither the global RNA pol II assembly defect observed in the *rtr1Δ* mutant cells (Figure S4A), nor the differences in chromatin-associated RNA pol II in relation to a wild-type strain (Figure S4B), in line with the incapability to suppress the sensitivity of the *rtr1Δ* mutant to 2% formamide phenotype (Figure 8A). Similar results were obtained by overexpressing *RPB7*, despite *RPB7* overexpression complementing the formamide sensitivity of the *rtr1Δ* mutant [27]. These results were expected, given the role of Rtr1 in RNA pol II biogenesis that involves different subunits, and not only Rpb4/7, probably in previous steps to Rpb4/7 association.

These data suggest that increasing the dose of the *RPB5* gene was enough to restore the assembly defect of RNA pol II provoked by lack of Rtr1 and the alteration to Rpb4 binding to chromatin-associated RNA pol II, leading to a partial suppression of the mRNA stability defect. These results indicate a direct role of Rpb4 in mRNA imprinting and decay in the *rtr1Δ* mutant cells. They also suggest additional mechanisms involving Rtr1, apart from the correct assembly of RNA pol II, operating for mRNA decay.

### Rtr1 cooperates with Rpb4 and Dhh1 for mRNA imprinting and chromatin association

Our above results revealed generally increased mRNA stability in *rtr1Δ* mutant cells in correlation with diminished Rpb4-mRNA imprinting, which likely resulted from a defect in Rpb4 binding to chromatin-associated RNA pol II, although additional mechanism involving Rtr1 seemed to act. Given the results describing that impairing Rpb4 association to mRNAs would lead to reduced mRNA decay [48,54] and that Rtr1 physically interacts with its own mRNA by autoregulating its turnover [44], we investigated the functional relation between Rtr1 and Rpb4, and analysed whether Rpb4 would also affect the association between Rtr1 and mRNA. To this end, we performed UV-crosslinking and mRNA isolation in the *rpb4Δ* mutant and its wild-type isogenic strains, both containing a functional Rtr1-TAP tagged version of this protein. Surprisingly, Rtr1 globally associated with mRNAs in both the wild-type and *rpb4Δ* mutant cells (see Figure 10A). Notably, *RPB4* deletion decreased the Rtr1 association with mRNAs, which indicates that lack of Rpb4 affects Rtr1-mRNA imprinting.

**Figure 10:**
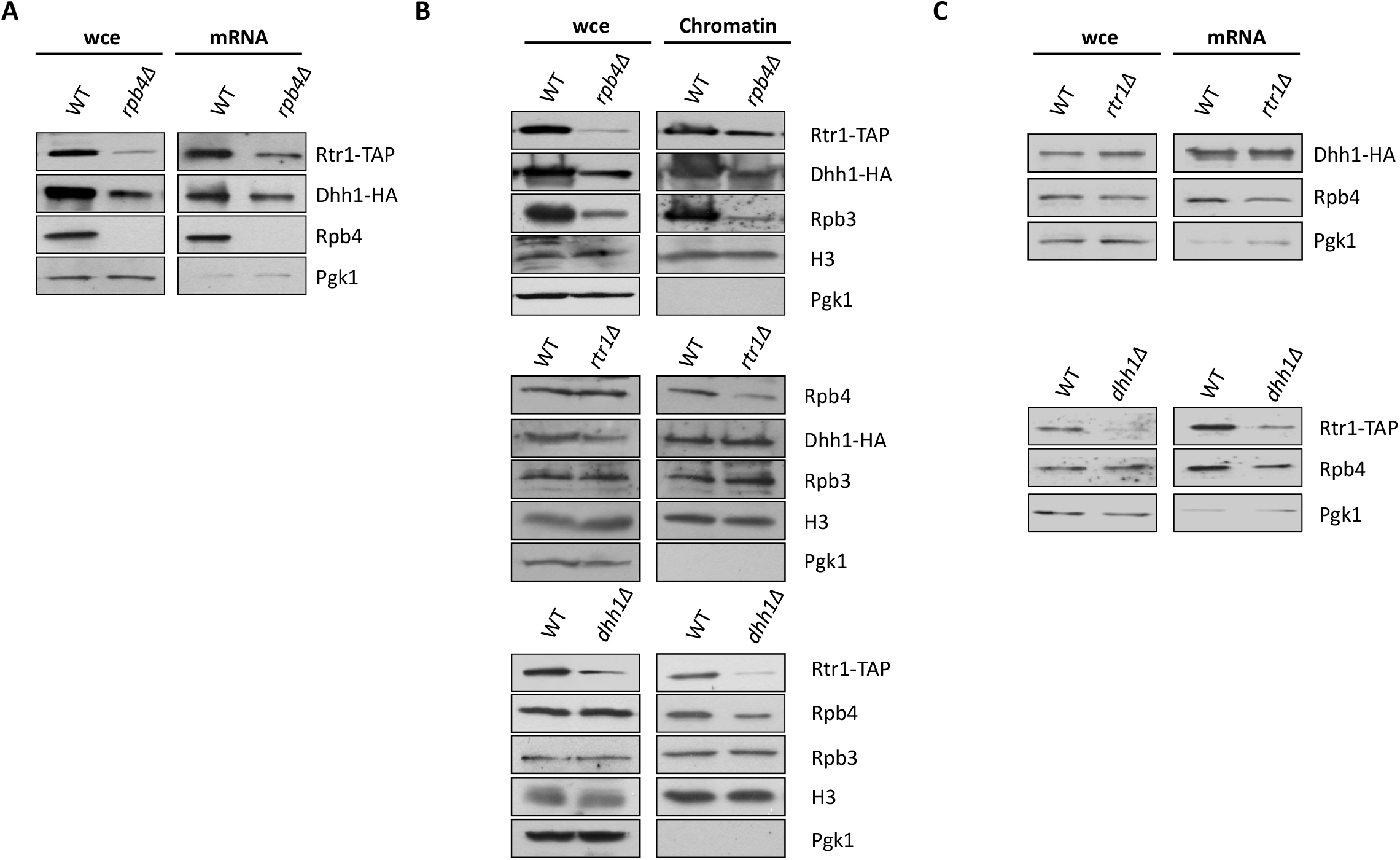
Rtr1, Rpb4 and Dhh1 cooperate to control mRNA stability. **A**) Western blot of the whole-cell crude extracts and oligo-dT purified mRNAs after exposure to 1,200 mJ/cm^2^ of 254 nm UV in the wild-type and *rpb4Δ* mutant grown in SD medium at 30°C. Pgk1 was used as a negative control of the mRNA-associated proteins. **B**) Western blot of the whole cell extracts and chromatin-enriched fractions obtained by the yChEFs procedure [58,62] from *rtr1Δ*, *rpb4Δ* and *dhh1Δ* mutants and their isogenic wild-type strains grown in YPD medium at 30°C. A similar amount of chromatin was purified from the wild-type and mutant strains as similar levels of histone H3 were detected. Pgk1 was used as a negative control of cytoplasmic contamination **C**) Western blot of the whole-cell crude extracts and oligo-dT purified mRNAs after exposure to 1,200 mJ/cm^2^ of 254 nm UV in the cells grown in SD medium at 30°C for *rtr1Δ* mutant and its wild-type strain (upper panel), and for *dhh1Δ* and its wild-type strain (lower panel). In A, B and C, Rpb4 was detected by a specific anti-Rpb4 antibody. Rtr1 and Dhh1 were detected using anti-TAP and anti-HA antibodies, respectively, in the strains containing an *rtr1::TAP* and/or *dhh1-HA* alleles. Note that chromatin and mRNA were properly isolated as no cytoplasmic Pgk1 protein was detected. All the experiments corresponded to at least three independent biological replicates.

The interaction of Rpb4 with mRNA occurs in the context of RNA pol II during transcription [48,55]. Moreover, Rtr1 has been described as an Ser5P-CTD phosphatase that participate in the transition from transcription initiation to elongation [29,31]. Thus we speculated that Rpb4 and Rtr1 could impact their association with chromatin and, consequently, mRNA imprinting. According to our data above, which revealed a reduction in the Rpb4 association with chromatin as a result of *RTR1* deletion, we also purified chromatin fractions by the yChEFs approach [58,62] from the Rtr1-TAP tagged *rpb4Δ* mutant and its wild-type isogenic strains, and we analysed the association of Rtr1 with chromatin by western blot (Figure 10B, top panel). We observed that lack of Rpb4 lowered the levels or Rtr1 associated with chromatin. It is worth noting that lack of Rpb4 seemed to also affect the global amount of cellular RNA pol II in our genetic background (see whole cell extracts, wce). Taken together, our data indicate that Rtr1 and Rpb4 cooperate to modulate their association with chromatin, and concomitantly, their mRNA imprinting.

It has been proposed that Rtr1 mediates the decay of its own mRNA by involving the RNA helicase Dhh1 and exonucleases Rex2 and Rex3 [44]. Furthermore, Dhh1 has been found to act as an interactor of Rtr1 [41]. By considering these data and the fact that lack of Dhh1 induces general mRNA stabilisation [71], we explored whether Dhh1 could also mediate the regulatory mechanisms of mRNA imprinting by involving Rtr1 and Rpb4. To do so, we first analysed the association of Dhh1 with mRNAs by performing UV-crosslinking experiments, as above, in the indicated *rpb4Δ* and *rtr1Δ* mutants and their isogenic wild-type strains, all containing Dhh1-HA tagged proteins. *RTR1* deletion did not affect the association Dhh1 with mRNAs, unlike what occurs for Rpb4 (Figure 10C, upper panel). In line with this, *RTR1* deletion did not alter Dhh1 association with chromatin (Figure 10B, middle panel). However, and notably, lack of Rpb4 decreased the association of Dhh1 with mRNAs, and similarly to what occurred for Rtr1 (Figure 10A). To gain more insight, we also analysed the association of Rpb4 and Rtr1 with mRNA by UV-crosslinking experiments in functional Rtr1-TAP tagged containing the *dhh1Δ* mutant and its isogenic wild-type strains. In line with the tripartite cooperation among these three proteins, lack of Dhh1 reduced the amount of Rtr1 and Rpb4 associated with mRNAs (Figure 10C, lower panel), and also with chromatin (Figure 10B, lower panel).

All these data collectively indicate that Rtr1, Rpb4 and Dhh1 cooperate to mediate mRNA stability and mRNA cotranscriptional imprinting.

## DISCUSSION

This work unravels the role of RNA pol II CTD Ser5-P phosphatase Rtr1 in the biogenesis of RNA pol II and in mRNA decay, and indicates that Rtr1 interconnects both processes. The effect of Rtr1 on RNA pol II biogenesis would favour its assembly by mediating the association of subassembly complexes to the rest of the enzyme, and would determinate the chromatin-associated RNA pol II population containing Rpb4 which would, in turn, cotranscriptionally impact mRNA decay. Our data also suggest that Rtr1 cooperates with Rpb4 and Dhh1 to mediate the mRNA degradation process.

Our results show a new role for Rtr1 in RNA pol II biogenesis, likely in the cytoplasm, by acting in the assembly of the enzyme, in addition to the previously proposed function for Rtr1 and its human ortholog RPAP2 as nuclear import factors involved in RNA pol II biogenesis [26,38]. As previously reported, the cytoplasmic assembly of RNA pol II is a sequential process, during which different subassembly complexes associate [14], by assuming the similar process described for bacterial RNA pol [72]. One of these, the Rpb1 subassembly, corresponds to subunits Rpb1, Rpb5, Rpb6 and Rpb8, and would appear to be associated with the rest on the enzyme in a final cytoplasmic RNA pol II assembly step [14]. Our results demonstrate that lack of Rtr1 affects the cytoplasmic assembly of RNA pol II by altering the association of Rpb5, Rpb6 and Rpb8, but also of dimer Rpb4/ Rpb7, with the rest of the enzyme, which suggests that Rtr1 mediates the correct assembly of the Rpb1 subassembly, likely in a final cytoplasmic step. In line with this role for Rtr1, our results reveal cytoplasmic Rpb1 accumulation under *RTR1* deletion, which agrees with previous observed results in yeast [26] and human cells under RPAP2 silencing [38]. In addition, a role of Rtr1 in the nuclear diffusion of small RNA pol II subunits has been previously proposed, and agrees with the minor cytoplasmic accumulation of some of them, such as Rpb3 and Rpb11, under *RTR1* deletion [26]. Although our results also demonstrate the cytoplasmic accumulation of Rpb4 by lack of Rtr1, we cannot speculate about its possible nuclear diffusion.

In line with the role of Rtr1 in RNA pol II assembly in human cells, RPAP2 has been found in association with small GTPases GPN1 and GPN3, GrinL1A, and R2TP/Prefoldin-like to cooperate in binding Rpb1 and in favouring its assembly in complete RNA pol II [15]. Accordingly in human cells treated with transcription inhibitor α-amanitin, which provokes Rpb1 degradation [73] and Rpb3 accumulation [15], RPAP2 has been found to be associated with the Rpb2 and Rpb3 subassembly complexes, which are assembled in an initial RNA pol II biogenesis step [14,15]. Furthermore, and in agreement with this role for Rtr1 mediating RNA pol II assembly, Npa3 and Gpn3 have been found to co-purify with Rtr1 [34,41] and Npa3 purification leads to the identification of a 10-subunit RNA Pol II lacking Rpb4/7 [34].

How the association of the Rpb4/Rpb7 subassembly complex occurs exactly is unknown [14]. However, we observed how the association of Rpb4/7 with the rest of the enzyme under *RTR1* deletion diminished, which suggests that it could assemble in a final cytoplasmic RNA pol II biogenesis step in cooperation with the Rpb1 subassembly complex, or even later, with the cooperation of Rtr1. We cannot rule out that the defect in Rpb6 association with the rest of the enzyme could also affect the association of Rpb4/7 by taking into account that Rpb6 contacts this dimer [4,63]. Nevertheless, our results do not rule out that Rtr1 is also involved in the nuclear association of Rpb4 with the rest of the enzyme. However, Rpb4 cytoplasmic accumulation in the cells lacking Rtr1 supports a major impact on Rpb4/7 cytoplasmic assembly in line with previous results showing no significant large population of free heterodimer Rpb4/7 in wild-type cells [34].

In agreement with a functional relation between Rtr1 and Rpb4/7, RNA pol II purification by using Rtr1-TAP as bait results in the isolation of a 10-subunit with the relative depletion of this dimer [34]. These results do not contradict those presented herein and could also support the notion that Rtr1 is necessary to favour Rpb4 (and likely Rpb7) assembly in the cytoplasm, or even in the nucleus. In fact Rtr1 interacts with both unmodified and phosphorylated RNA pol II in CTD-Ser5 [27,29]. Similarly, *RTR1* shows genetic interactions with *RPB4* and *RPB7* ([27] and our results), and *rtr1Δ* genetically interacts with the *rpb1* and *rpb6* mutants that affect integrity or RNA pol II, and lead to the dissociation of either dimer Rpb4 or Rpb4/7 ([63], and our data).

Lack of Rtr1 leads to the nuclear accumulation of two populations of chromatin-associated RNA pol II, complete or lacking Rpb4, that are likely functional. Furthermore, other intermediary complexes or free subunits can also accumulate, as our results show for Rpb1 and Rpb4, or in line with previously reported findings [26]. These subcomplexes or free subunits, if they shuttle to the nucleus, do not seem to be associated with chromatin. So we cannot rule out that some assembly steps also occur in the nucleus if we bear in mind previous propositions [16,26]. It has been proposed that Rtr1 and other elongation factors may influence Rpb4/7 dissociation with the rest of RNA pol II once interacting during transcription [34], although the occurrence of Rpb4/7 dissociation is controversial [34,65,66]. Our ChIP data do not indicate any major contribution of Rtr1 to this process that could account for the decrease in Rpb4 association with the rest of the enzyme provoked by lack or Rtr1. However, we cannot rule out that this phenomenon may occur for a specific group of genes in line with a previous proposition [34].

Our data also depict Rtr1 as specific for RNA pol II assembly because *RTR1* deletion did not seem to affect RNA pol I and III assembly. However, Rtr1 seems to cooperate with Bud27, a prefoldin-like that mediates the assembly of the three RNA pols in an Rpb5-dependent manner [22]. The data herein shown suggest that Bud27 could participate by favouring Rpb1 subassembly complex formation and association, likely in cooperation with Rtr1, if we take into account that Bud27 allows the correct cytoplasmic assembly of Rpb5 and Rpb6 to the rest of the enzyme before its nuclear entry [22]. In line with this, *RPB5* overexpression overcomes the RNA pol II assembly defects of *rtr1Δ* cells. Accordingly, the Bud27 human counterpart, URI, has been found in a subassembly intermediary containing RPB5 [15,16,22,24].

Previous results have demonstrated a role for Rpb4 (and also of dimer Rpb4/7) in the life cycle of mRNA by imprinting it cotranscriptionally, and later accompanying it during the export to the cytoplasm, translation, mRNA decay or cytoplasmic accumulation. Lack of Rpb4 increases mRNA stability [48,53,54,55,56,74]. Similarly, increased mRNA stability by *RTR1* deletion was observed herein. It is worth noting that *RTR1* deletion alters the association of Rpb4 with RNA pol II. So we propose as a first option that increased mRNA stability could be the consequence of altering the RNA pol II assembly and Rpb4 binding to chromatin-associated RNA pol II as it is also accompanied by reduced Rpb4-mRNA imprinting. In line with this, mutations that alter the RNA pol II assembly and provoke the dissociation of Rpb4 from the rest of the complex also increase mRNA stability by decreasing Rpb4-mRNA imprinting [48,53,55,63]. Our data show that *RPB5* overexpression overcomes the alteration to Rpb4 binding to the chromatin-associated RNA pol II in *rtr1Δ* cells and partially suppresses mRNA stability alteration, which denotes a direct role of Rpb4-bound to chromatin-associated RNA pol II in mRNA decay in the cells deleted for *RTR1*. In line with this, *RPB4/7* overexpression has been shown to also partially suppress the mRNA stability increase provoked by the *rpb6^Q100R^* mutation that affects the Rpb4/7 association with the rest of the enzyme [55]. Our data also point out that Rpb4-bound to chromatin-associated RNA pol II is a key element in Rpb4-mRNA imprinting, which coincides with previous propositions [74], as lack of Rtr1 does not alter the global Rpb4 amount, but increases the fraction of the free Rpb4 subunit. The fact that *RPB4/7* overexpression did not overcome either the RNA pol II assembly of *rtr1Δ* cells, or formamide sensitivity, which occurs by *RPB5* overexpression, this agrees with a major role of Rtr1 in the cytoplasmic assembly of the Rpb1 subassembly complex and argues for later Rpb4/7 association in line with the previously proposed RNA pol II assembly model [14].

A second option, that must act in addition to the first one, is that Rtr1 plays a specific role in mRNA decay, which falls in line with previous data reported for this protein autoregulating the decay of its own mRNA [44]. Strikingly our results suggested that Rtr1 may imprint a broad population of mRNAs. Rtr1-mRNA imprinting may occur in cooperation with Rpb4, and likely cotranscriptionally, as lack of this RNA pol II subunit reduces the amount of Rtr1 associated with chromatin and also decreases Rtr1-mRNA imprinting. These results hint that Rpb4 cotranscriptionally modulates Rtr1-mRNA imprinting. This phenomenon may occur during the transition from transcription initiation to elongation if we take into account the role of Rtr1 as a Ser5-P phosphatase [29,31] and the involvement of Rpb4/7 in the RNA pol II conformational change that concomitantly occurs not only in yeast, but also in other organisms [75,76,77]. However, we cannot rule out that this imprinting occurs later during transcription elongation as Rtr1 occupies the whole gene body during transcription and coincides with Ser5P, but also with Ser2P [29]. Coinciding with this possibility, Rtr1 physically interacts with different transcription elongation factors, such as Dst1, Spt5, Rba50, or with some subunits of the PAF complex [41]. In any case, Rtr1-mRNA imprinting may occur during the transition from transcription elongation to termination if we bear in mind the proposed role of Rtr1 in transcription termination [36], and also as CTD Tyr1-P phosphatase [31].

A previous functional cooperation between Rtr1 and Dhh1 in *RTR1* mRNA decay regulation has been proposed, which also involves Rex exonucleases [44]. Our results reveal reduced Dhh1 chromatin association when Rpb4 is lacking, which agrees with data previously reported for Dhh1 associated with gene promoters [47]. This reduction is accompanied by reduced Dhh1-mRNA imprinting, which suggests that Dhh1 imprints mRNA cotranscriptionally in cooperation with Rpb4. Accordingly, these results suggest a defect in mRNA decapping provoked by lack of Rpb4. However, no major effects on the Dhh1 association with chromatin or Dhh1-mRNA imprinting occur when Rtr1 is lacking, despite the described cooperation between these proteins [44]. It is worth noting that *DHH1* deletion affects the association of Rtr1 with both chromatin and mRNA. Accordingly, we speculate that Dhh1-Rtr1 cooperation for mRNA decay regulation may occur mainly in the cytoplasm. Alternatively, Rtr1 could impact Dhh1 chromatin and mRNA association, albeit weakly. It is worth noting that Dhh1 has been found to interact with Rtr1 in proteomic studies [41].

Taking together, we propose that the functional cooperation of Rtr1, Rpb4 and Dhh1 could operate to modulate mRNA decay by then connecting transcriptional and mRNA degradation machinery as elements of the crosstalk between mRNA synthesis and decay [47]. Our results herein described and those of other authors allow us to hypothesize the model shown in Figure 11. Rtr1 acts in RNA pol II assembly to allow the correct cytoplasmic assembly of subassembly complexes Rpb1 and Rpb4/7 in a final RNA pol II biogenesis step. The whole RNA pol II would later be imported to the nucleus. Furthermore, we cannot rule out that Rtr1 favours Rpb4 association with the rest of the enzyme in the nucleus. However, lack of Rtr1 would affect RNA pol II assembly and lead to obtain both the complete enzyme and a RNA pol II lacking Rpb4, which would be associated with chromatin, as well as other intermediary complexes and free subunits. During transcription, Rtr1 would associate with RNA pol II (and, thus, with chromatin) in an Rpb4-dependent manner. Rpb4 would cooperate with Rtr1 for Rtr1-mRNA imprinting, but would also imprint mRNA, and both would do so cotranscriptionally. Nor can we rule out a mutual cooperation between Rtr1 and Rpb4 for mRNA imprinting in line with Rpb4 cooperating with RBP Puf3 to imprint and modulate mRNA stability for a subset of genes [74]. Similarly, Rpb4 would influence both the association of Dhh1 with chromatin and Dhh1 cotranscriptional mRNA imprinting. Triple Rtr1-Rpb4-Dhh1 mRNA imprinting would allow the correct mRNA decay regulation in the cytoplasm by the action of 3’ and 5’ polyA-mRNA degradation machinery. Under *RTR1* deletion, mRNA imprinting would be altered and affect cytoplasmic mRNA decay, which would consequently increase mRNA stability.

**Figure 11:**
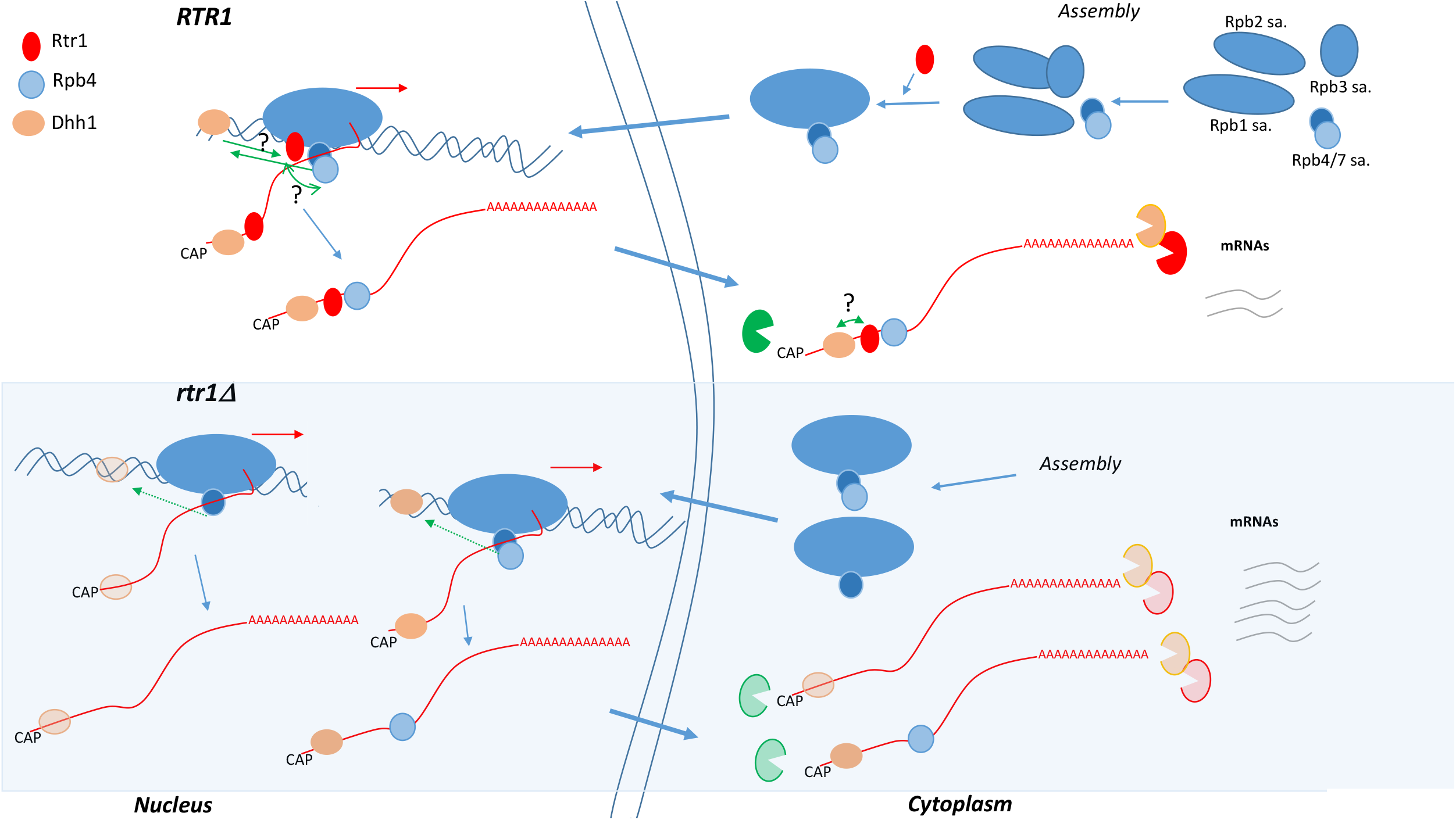
Model for the Rtr1 role in RNA pol II cytoplasmic assembly and for Rtr1, Rpb4 and Dhh1 cooperation in mRNA decay. In the cytoplasm, Rtr1 would act in RNA pol II assembly to facilitate the association of Rpb1 and Rpb4/7 subassembly complexes in the final RNA pol II biogenesis steps. Complete RNA pol II, or that lacking Rpb4 (under *RTR1* deletion), would be transported to the nucleus, likely by an Iwr1-dependent pathway. We cannot rule out that Rtr1 also acts by promoting the nuclear association of Rpb4 with the rest of the enzyme. In the nucleus, Rtr1 would associate with RNA pol II and would cotranscriptionally imprint mRNA in cooperation with Rpb4, which also imprints mRNA cotranscriptionally. Rpb4 would also cooperate with Dhh1 for chromatin association and cotranscriptional mRNA imprinting. *RTR1* deletion would affect RNA pol II biogenesis and Rpb4 association with the rest of the enzyme, as well as the transcription mediated by RNA pol II, and Rtr1- and Rpb4-mRNA imprinting. Furthermore, RNA pol II lacking Rpb4 would affect Dhh1 chromatin association and Dhh1-mRNA imprinting. Consequently, lack of Rtr1 would lead to altered imprinted mRNAs, which would be exported to the cytoplasm that would increase their mRNA stability by altering the actions of 3’ and 5’ mRNA degradation machinery.

Our data suggest Rtr1 regulating the decay of a large mRNA population instead of only its own mRNA [44]. Consequently, it would be interesting to define this Rtr1-associated mRNA population. The possibility of this matching at least part of the Rpb4 one, as is the case for Rpb4 and RBP Puf3 [74], would allow us to further analyse the specific role of Rtr1 in mRNA decay regulation in detail. Finally, investigating whether other RNA pol II CTD phosphatases and kinases would also be involved in mRNA decay would involve including new layers in the crosstalk between mRNA synthesis and decay.

## MATERIALS AND METHODS

### Yeast strains, genetic manipulations, media and genetic analysis

The common yeast media, growth conditions and genetic techniques were used as described elsewhere [78]. Formamide sensitivity was tested using a 2% dilution as previously described [27]. Yeast strains and plasmids are listed in Supplementary Tables S1 and S2, respectively.

The strains containing an *rtr1Δ::KanMX4* allele in our working yeast backgrounds, except YFN562, were obtained by chromosomal integration of a PCR product amplified using genomic DNA from strain YFN161 containing the *rtr1Δ::KanMX4* construction as templates (haploid strain derived from the Y26137 diploid strain, EUROSCARF) with oligonucleotides Rtr1-501 and Rtr1-301 (Table S3). The YFN556 strain containing an *rtr1Δ::kanMX4::HIS3* allele was obtained by integrating the *HIS3* maker from plasmid M4754 into the *rtr1Δ::KanMX4* marker of the YFN160 strain by chromosomal integration. Other strains containing the *rtr1Δ::kanMX4::HIS3* allele in our working yeast backgrounds were obtained by the chromosomal integration of a PCR product amplified using genomic DNA from strain YFN556 with oligonucleotides Rtr1-501 and Rtr1-301 (Table S3) *rtr1::TAP::HIS3MX6* in our working yeast backgrounds, except for the YFN415 strain, resulted from chromosomal integration of a PCR product amplified using genomic DNA from a strain containing *rtr1::TAP::HIS3MX6* (OpenByosystem) with oligonucleotides Rtr1-501 and Rtr1-301, respectively, as templates (Table S3).

The strains bearing the derived plasmids encoding *RPB5* alleles were generated by plasmid shuffling of *pFL44-RPB5* (2 μm, URA3) [79].

### Protein extract preparation and immunoprecipitation

Protein whole cell extracts and protein immunoprecipitation were performed as described [63]. Briefly for protein extract preparation, 150 ml of cells growing exponentially (OD_600_ 0.6-0.8) were centrifuged, pelleted and resuspended in 0.3 ml of lysis buffer (50 mM HEPES [pH 7.5], 120 mM NaCl, 1 mM EDTA, 0.3% 3-[(3-cholamidopropyl)-dimethylammonio]-1-propanesulfonate (CHAPS)) supplemented with 1X protease inhibitor cocktail (Complete; Roche), 0.5 mM phenylmethylsulfonyl fluoride (PMSF), 2 mM sodium orthovanadate, and 1 mM sodium fluoride. Cell disruption was carried out by vortexing (3 cycles, 5 min each) at 4°C using 0.2 ml of glass beads (425-600 μm; Sigma). For the Rpb3 immunopurification, 1μg of anti-Rpb3 antibody (anti-POLR2C;1Y26, Abcam) was coupled to 50 μl of Dynabeads Sheep-anti-Mouse IgG (Invitrogen) per sample, and 2 mg of a whole cell protein extract were used for each immunoprecipitation. A similar procedure was followed for TAP pull down, with 50 μl of Dynabeads-Pan Mouse (Invitrogen). The affinity-purified proteins were released from the beads by boiling for 10 min and were analysed by western blotting with different antibodies.

### SDS-PAGE and western blot analysis

Protein electrophoresis and western blot were carried out as described in [63].

For the western blot analyses, anti-CTD [58], anti-Rpb3 (anti-POLR2C;1Y26, Abcam), anti-Rpb4 (Pol II RPB4 (2Y14); Biolegend), anti-Rpb5 (a polyclonal antibody generated against *S. cerevisiae* Rpb5 in our lab), anti-Rpb6 (a gift from M. Werner), anti-Rpb7 (Rpb7 (yN-19); Santa Cruz Biotechnology), anti-phosphoglycerate kinase, Pgk1 (22C5D8; Invitrogen), anti-H3 (ab1791; Abcam), anti-hemaglutinin (anti-HA; 12CA5; Roche) and PAP (*Sigma*) antibodies were used.

### Fluorescence Microscopy

The cells expressing Rpb1-Gfp and Gfp-Rpb4 were grown at 30°C in SD medium supplemented with the corresponding amino acids to reach an OD_600_ ~ 0.5-0.7. For Rpb4 localization, strains were transformed with the centromeric vector expressing a Gfp-Rpb4 fusion protein [56] (Supplementary Table S2). Slides were covered with Vectashield mounting solution (Vector Laboratories). Fluorescence intensity was scored with a fluorescence microscope (Olympus BX51).

### Yeast Chromatin-Enriched Fractions preparation

Chromatin-enriched fractions preparation was carried out by the yChEFs procedure [58,62] using 75 ml of the YPD cultures grown exponentially (OD_600_ ~0.6– 0.8). The chromatin-bound proteins were resuspended in 1X SDS-PAGE sample buffer, boiled for 10 min and analysed by western blotting with different antibodies.

### Isolation of the mRNA-associated proteins

mRNA crosslinking was carried out as described elsewhere [48] with some modifications. Briefly, 250 ml of cell cultures grown in SD medium (with requirements) to an OD_600_ ~ 0.6-0.8 were exposed to 1,200 mJ/cm^2^ of 254 nm UV in a UV crosslinker (Biolink Shortwave 254 nm), resuspended in 350 μl of lysis buffer (20 mM Tris pH 7.5, 0.5 M o NaCl, 1 mM EDTA, 1X protease inhibitor cocktail [Complete; Roche]) and 200 μl of glass beads (425-600 μm, Sigma) and broken by vortexing for 15 min at 4°C. An aliquot of lysate was used as the INPUT control. Lysate was incubated with 150 μl of oligo (dT)_25_ cellulose beads (New England BioLabs, cat no. S1408S) for 15 min at room temperature. PolyA-containing RNA isolation and elution were carried out as previously described [48]. The mRNA-associated proteins were analysed by SDS-PAGE and western blot with the appropriate antibodies.

### mRNA stability analysis

mRNA stability analysis was performed as previously described [48]. Briefly, 150 ml of cells were grown in SD (with requirements) to reach an OD_600_ ~ 0.5. Cells were treated with 5 μg/ml of thiolutin. Next 15-ml aliquots of cell samples were taken at different times after thiolutin addition (up to 100 min), pelleted and frozen. Total RNA was isolated from these samples and cDNA was synthesized as previously described [63]. The mRNA stability (half-lives, HL) for the selected genes was analysed following the decay curves obtained by RT-qPCR with specific primers for the corresponding genes (see Table S3).

For total mRNA half-live calculation, the previously obtained RNA was used for dot-blot analysis: 2 μg of total RNA dots were added to a nitrocellulose membrane previously washed with SSC 2X buffer (SSC 20X: NaCl 3M, sodium citrate 300 mM, pH 7). The membrane was incubated for 5 min at 65°C and exposed to 400 mJ/cm^2^ of 254 nm UV in a UV crosslinker (Biolink Shortwave 254 nm). After crosslinking, the membrane was incubated in prewarmed hybridisation buffer (NaPO4 0.5 M, pH 7, EDTA 10 mM, SDS 7%) at 56°C for 45-60 min. Then 50 pmol of oligodTCy3 were added and incubated overnight at the same temperature. The membrane was washed 6 times at room temperature using prewarmed washing buffer (NaPO4 0.28 M, pH 7.2, SDS 7%). Fluorescence was analysed with a Molecular Imager^®^ VersaDoc™ MP system at 635 nm and the Quantity one software. The graphical quantifications of the dot blot are shown and represented on a natural logarithmic scale.

The analysis of *GAL1* and *GAL10* genes mRNA stability was performed as previously described [48] by shifting the cells that grew exponentially (OD_600_ ~ 0.5-0.6) from SD-galactose to SD-glucose to stop transcription. Cell samples were collected at different time points after glucose addition (5, 10, 15, 20 and 30 min). RNA extraction was performed as described above. mRNA stability was analysed by RT-qPCR with specific primers for the corresponding genes (see Table S3).

### Quantitative real-time PCR (RT-qPCR)

Real-time PCR was performed in a CFX-384 Real-Time PCR instrument (BioRad) with the EvaGreen detection system “SsoFast™ EvaGreen^®^ Supermix” (BioRad). Reactions were performed using cDNA corresponding to 0.1 ng of total RNA in 5 μl of total volume. Each PCR reaction was performed at least 3 times with three independent biological replicates to obtain a representative average. The 18S rRNA gene was used as a normalizer. The employed oligonucleotides are listed in Supplementary Table S3.

### Chromatin immunoprecipitation

The chromatin immunoprecipitation experiments were performed using anti-Rpb3 (anti-POLR2C;1Y26, Abcam) or anti-Rpb4 (Pol II RPB4 (2Y14); Biolegend) as previously described [63]. For real-time PCR, a 1:100 dilution was employed for the input DNA and a 1:4 dilution for the immunoprecipitated samples DNA. Genes were analysed by quantitative real-time PCR in a CFX-384 Real-Time PCR instrument (BioRad) in triplicate with at least three independent biological replicates using SYBR premix EX Taq (Takara). Quantification was performed as indicated in the Figure legends. The oligonucleotides utilized for the different PCRs are listed in Supplemental Table S3.

## ACKNOWLEDGEMENTS

We thank Dr. J.E. Pérez-Ortín and Dr. Olga Calvo for their helpful discussion and the “Servicios Centrales de Apoyo a la Investigación (SCAI)” of the University of Jaen for technical support.

