## Supplemental Material for "RNA polymerase II assembly and mRNA decay regulation are mediated and interconnected *via* CTD Ser5P phosphatase Rtr1 in *Saccharomyces cerevisiae*"

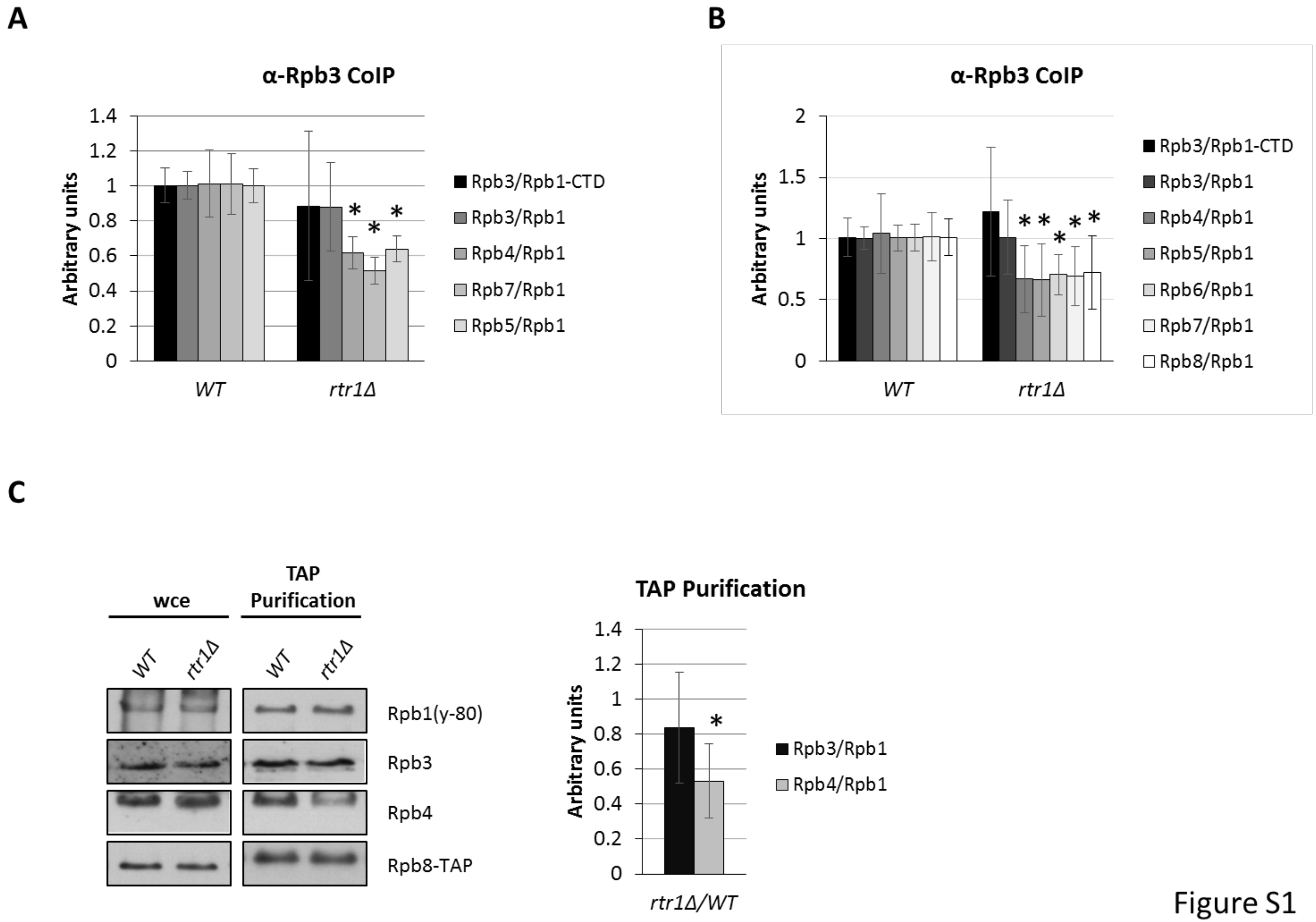


Figure S1

**Figure S1: *RTR1* deletion affects the assembly of RNA pol II. A** and **B** correspond to the quantification of the western blot from the RNA pol II immunoprecipitated from the *rtr1Δ* mutant and wild-type strains, from Figure 1A and B, respectively, showing the ratio for each subunit *vs.* Rpb1 (y-80). **C**) The quantification of the western blot from Rpb8-TAP pulldown showing the ratios of Rpb3 and Rpb4 RNA pol II subunits *vs.* Rpb1 (y-80). *$p<0.05$ (t-test).

A

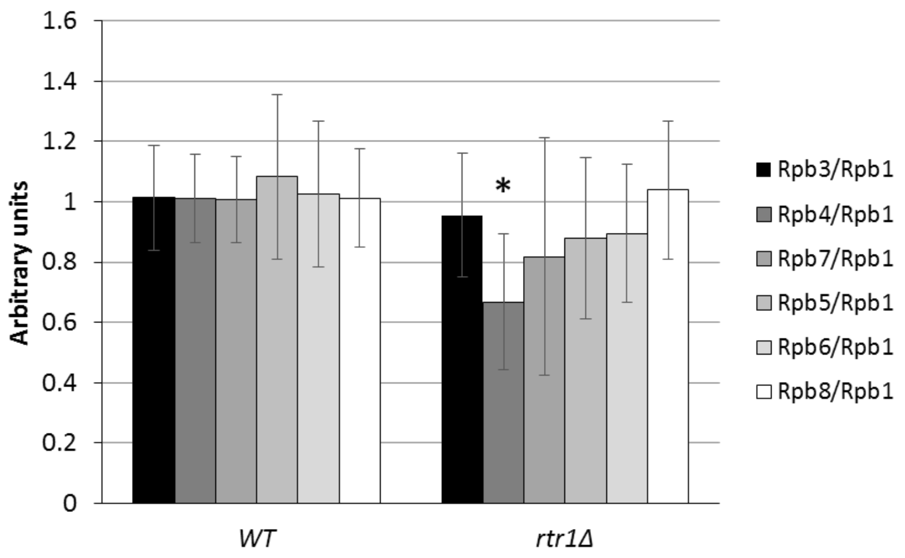


Figure S2

**Figure S2: *RTR1* deletion affects the assembly of RNA pol II.** The quantification of the western blot for different RNA pol II subunits *vs*. Rpb1 from chromatin purification, corresponding to the results in Figure 4 for the *rtr1Δ* mutant and wild-type strains. *p<0.05 (t-test).

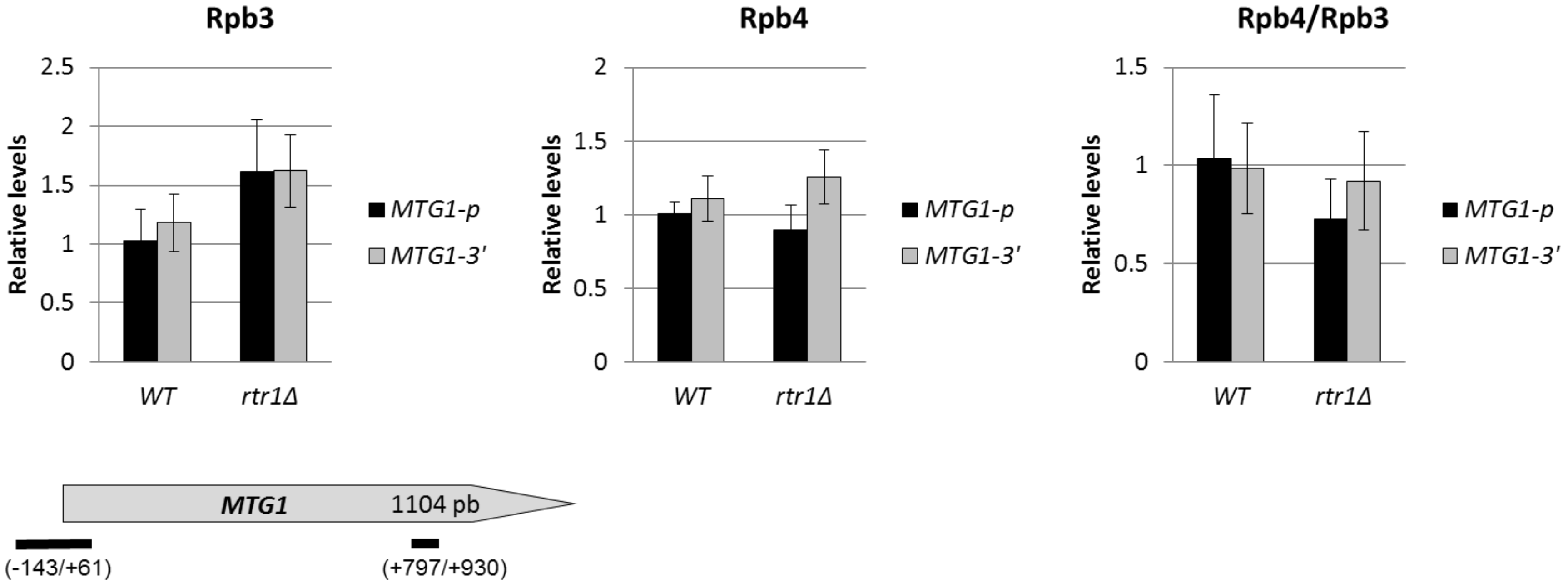


Figure S3

**Figure S3: The *rtr1Δ* mutation affects gene occupancy by RNA pol II, but not global Rpb4 dissociation.** Chromatin immunoprecipitation (ChIP) analysis for the *MTG1* gene in the wild-type and *rtr1Δ* cells performed with anti-Rpb3 (left panel) and anti-Rpb4 (middle panel) antibodies, against Rpb3 and Rpb4 RNA pol II subunits. Right panel: the Rpb4/Rpb3 ratios for the Rpb4 dissociation analysis, from the left and middle panel's results. Lower panel: the transcription unit used in this work indicating the location of the analysed PCR amplicons. The values found for the immunoprecipitated PCR products were compared to those of the total input, and the ratio of each PCR product of the transcribed genes to a non-transcribed region of chromosome V was calculated.

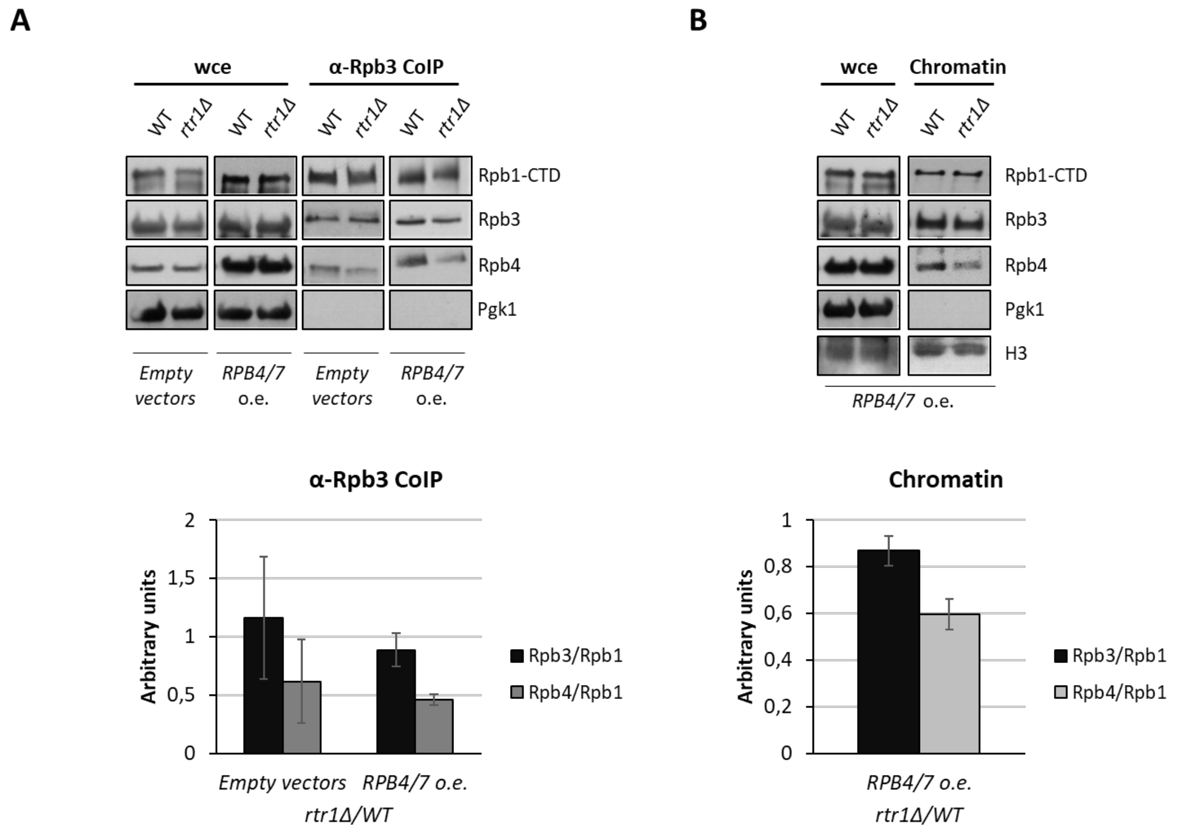


Figure S4

**Figure S4: RPB4/7 overexpression does not overcome the RNA pol II assembly defect in the *rtr1Δ* mutant. A**) Rpb3 immunoprecipitation in the wild-type and *rtr1Δ* cells both overexpressing the *RPB4/7* genes from high copy number plasmids or harbouring the corresponding empty vectors, grown in SD medium at 30ºC. The western blots of Rpb1, Rpb3 and Rpb4, and Pgk1 from the whole cell crude extracts and immunoprecipitated samples are shown (upper panel). Lower panel: the quantification of the western blot showing the *rtr1Δ* mutant/wild-type strains ratio for each subunit *vs.* Rpb1. Median and SD of two independent biological replicates. **B**) The whole-cell extract and chromatin enriched fractions obtained by the yChEFs procedure [1,2] from the same strains used in **A** overexpressing *RPB4/7* genes from high copy number plasmids. Cells were grown in SD medium at 30ºC and proteins were analysed by western blot with specific antibodies against Rpb1, Rpb3 and Rpb4. H3 histone was used as a positive control of the chromatin-associated protein and Pgk1 employed as a negative control of cytoplasmic contamination. Lower panel: the quantification for each RNA pol II subunit *vs.* Rpb1. Median and SD of two independent biological replicates.

**Supplementary Table S1.** ***Saccharomyces cerevisiae*** **strains**

| **Strain** | **Genotype** | **Origin** |
|---|---|---|
| BY4741 | *MATa his3-Δ1 leu2-Δ0 met15-Δ0 ura3-Δ0* | Euroscarf |
| Y26137 | *MATa/MATα his3-Δ1/his3-Δ1 leu2-Δ0/leu2-Δ0 lys2-Δ0/LYS2 MET15/met15-Δ0 ura3-Δ0/ura3-Δ0 rtr1Δ::kanMX4/RTR1* | Euroscarf |
| YFN160 | *MATα his3-Δ1 leu2-Δ0 ura3-Δ0 rtr1Δ::KanMX4* | This work |
| YFN161 | *MATa his3-Δ1 leu2-Δ0 lys2-Δ0 ura3-Δ0 rtr1Δ::KanMX4* | This work |
| YFN556 | *MATα his3-Δ1 leu2-Δ0 ura3-Δ0 rtr1Δ::kanMX4::HIS3* | This work |
| Y07202 | *MATa his3-Δ1 leu2-Δ0 met15-Δ0 ura3-Δ0 trp1Δ::kanMX4* | Euroscarf |
| YFN756 | *MATa his3-Δ1 leu2-Δ0 met15-Δ0 ura3-Δ0 trp1Δ::kanMX4 rtr1Δ::kanMX4::HIS3* | This work |
| YFN416 | *MATa his3-Δ1 leu2-Δ0 met15-Δ0 ura3-Δ0 rpb1::GFP::HIS5* | [3] |
| YFN744 | *MATa his3-Δ1 leu2-Δ0 met15-Δ0 ura3-Δ0 rpb1::GFP::HIS5 rtr1Δ::KanMX4* | This work |
| Rpb8-TAP | *MATa his3-Δ1 leu2-Δ0 met15-Δ0 ura3-Δ0 rpb8::TAP::HIS3Mx6* | Open Biosystems |
| YFN760 | *MATa his3-Δ1 leu2-Δ0 met15-Δ0 ura3-Δ0 rpb8::TAP::HIS3Mx6 rtr1Δ::KanMX4* | This work |
| YFN223 | *MATα ade2-101 his3-Δ200 leu2-Δ1 lys2-801 trp1-Δ63 ura3-52 YOR341W(RPA190)::3HA::HIS3 YOR116C(RPC160)::13Myc::TRP* | [3] |
| YFN743 | *MATα ade2-101 his3-Δ200 leu2-Δ1 lys2-801 trp1-Δ63 ura3-52 YOR341W(RPA190)::3HA::HIS3 YOR116C(RPC160)::13Myc::TRP rtr1Δ::KanMX4* | This work |
| YFN106 | *MATα his3-Δ1 leu2-3,112 lys2-801 met15-Δ0 trp1-Δ63 ura3-Δ0 bud27Δ::KanMX4* | [3] |
| Rtr1-TAP | *MATa his3-Δ1 leu2-Δ0 met15-Δ0 ura3-Δ0 rtr1::TAP::HIS3Mx6* | Open Biosystems |

| | | |
|---|---|---|
| YFN415 | *MATα his3-Δ1 Leu2* lys2-801 met15-Δ0 trp1-Δ63 ura3-Δ0 rtr1::TAP::HIS3Mx6 bud27Δ::KanMX4* | This work |
| Bud27-TAP | *MATa his3-Δ1 leu2-Δ0 met15-Δ0 ura3-Δ0 bud27::TAP::HIS3Mx6* | Open Biosystems |
| YFN742 | *MATa his3-Δ1 leu2-Δ0 met15-Δ0 ura3-Δ0 bud27::TAP::HIS3Mx6 rtr1Δ::KanMX4* | This work |
| Y01285 | *MATa his3-Δ1 leu2-Δ0 met15-Δ0 ura3-Δ0 rpb4Δ::KanMX4* | Euroscarf |
| Y03858 | *MATa his3-Δ1 leu2-Δ0 met15-Δ0 ura3-Δ0 dhh1Δ::KanMX4* | Euroscarf |
| PAY749 | *MATa his3-Δ1 leu2-Δ0 met15-Δ0 ura3-Δ0 dhh1::HA::HIS3* | Gift from Paula Alepuz |
| YFN736 | *MATa his3-Δ1 leu2-Δ0 met15-Δ0 ura3-Δ0 rtr1Δ::kanMX4::HIS3 dhh1Δ::KanMX4* | This work |
| YFN738 | *MATa his3-Δ1 leu2-Δ0 met15-Δ0 ura3-Δ0 dhh1Δ::KanMX4 rtr1::TAP::HIS3MX6* | This work |
| YFN739 | *MATa his3-Δ1 leu2-Δ0 met15-Δ0 ura3-Δ0 rtr1Δ::KanMX4 dhh1::HA::HIS3* | This work |
| YFN740 | *MATa his3-Δ1 leu2-Δ0 met15-Δ0 ura3-Δ0 dhh1::HA::HIS3 rtr1::TAP::HIS3MX6* | This work |
| YFN741 | *MATa his3-Δ1 leu2-Δ0 met15-Δ0 ura3-Δ0 dhh1::HA::HIS3 rtr1::TAP::HIS3MX6 rpb4Δ::KanMX4* | This work |
| YFN562 | *MATa his3-Δ1 leu2-Δ0 ura3-Δ0 bud27Δ::KanMX4 rtr1Δ::kanMX4::HIS3* | This work |
| YFN116 | *MATα his3-Δ200 leu2-3,112 trp1-Δ63 ura3-52 rpb1-Δ187::HIS3 + pYEB220 (2µm LEU2 RPB1)* | [4] |
| YFN117 | *MATα his3-Δ200 leu2-3,112 trp1-Δ63 ura3-52 rpb1-Δ187::HIS3 + pYEB220-rpo21-4 (2µm LEU2 rpo21-4)* | [4] |
| W303 | *MATa ade2-1 can1-100 his3-11,15 leu2-3,112 trp1-1 ura3-1* | [5] |
| OCSC2115 | *MATα ade2-1 can1-100 his3-11,15 leu2-3,112 trp1-1 ura3-1 rpb4Δ::HIS3* | Gift from Olga Calvo |
| YFN567 | *MATa ade2-1 can1-100 his3-11,15 leu2-3,112 trp1-1 ura3-1 rtr1Δ::KanMX4* | This work |

| | | |
|---|---|---|
| YFN638 | *MATa ade2-1 can1-100 his3-11,15 leu2-3,112 trp1-1 ura3-1 rpb4Δ::HIS3 rtr1Δ::KanMX4* | This work |
| YFN2 | *MATa ade2-101 his3-Δ200 leu2-Δ1 lys2-801a trp1-Δ63 ura3-52 rpb5Δ::ura3::LEU2 + pFL44L-RPB5 (2 μm URA3 RPB5)* | [6] |
| YFN750 | *MATa ade2-101 his3-Δ200 leu2-Δ1 lys2-801a trp1-Δ63 ura3-52 rpb5Δ::ura3::LEU2 + pFL44L-RPB5 (2 μm URA3 RPB5) rtr1Δ::KanMX4* | This work |
| YFN794 | *MATa ade2-101 his3-Δ200 leu2-Δ1 lys2-801a trp1-Δ63 ura3-52 rpb5Δ::ura3::LEU2 + + pASZ11-RPB5 (CEN ADE2 RPB5)* | This work |
| YFN789 | *MATa ade2-101 his3-Δ200 leu2-Δ1 lys2-801a trp1-Δ63 ura3-52 rpb5Δ::ura3::LEU2 + + pASZ11-RPB5 (CEN ADE2 RPB5) rtr1Δ::KanMX4* | This work |
| YFN5 | *MATa ade2-101 his3-Δ200 leu2-Δ1 lys2-801a trp1-Δ63 ura3-52 rpb5Δ::ura3::LEU2 + pGEN-rpb5-P151T (2μm TRP1 rpb5-P151T)* | [6] |
| YFN6 | *MATa ade2-101 his3-Δ200 leu2-Δ1 lys2-801a trp1-Δ63 ura3-52 rpb5Δ::ura3::LEU2 + pGEN-rpb5-H147R (2μm TRP1 rpb5-H147R)* | [6] |
| YFN49 | *MATa ade2-101 his3-Δ200 leu2-Δ1 lys2-801a trp1-Δ63 ura3-52 rpb5Δ::ura3::LEU2 + pGEN-rpb5-R200E (2μm TRP1 rpb5-R200E)* | [7] |
| YFN790 | *MATa ade2-101 his3-Δ200 leu2-Δ1 lys2-801a trp1-Δ63 ura3-52 rpb5Δ::ura3::LEU2 + pGEN-RPB5 (2μm TRP1 RPB5) rtr1Δ::KanMX4* | This work |
| YFN791 | *MATa ade2-101 his3-Δ200 leu2-Δ1 lys2-801a trp1-Δ63 ura3-52 rpb5Δ::ura3::LEU2 + pGEN-rpb5-P151T (2μm TRP1 rpb5-P151T) rtr1Δ::KanMX4* | This work |
| YFN792 | *MATa ade2-101 his3-Δ200 leu2-Δ1 lys2-801a trp1-Δ63 ura3-52 rpb5Δ::ura3::LEU2 + pGEN-rpb5-H147R (2μm TRP1 rpb5-H147R) rtr1Δ::KanMX4* | This work |

| | | |
|---|---|---|
| YFN793 | *MATa ade2-101 his3-Δ200 leu2-Δ1 lys2-801a trp1-Δ63 ura3-52 rpb5Δ::ura3::LEU2 + pGEN-rpb5-R200E (2µm TRP1 rpb5-R200E) rtr1Δ::KanMX4* | This work |
| WY204 | *MATα ade2 his3-Δ200 leu2-3,112 lysΔ201 ura3-52 rpb6Δ::HIS3 + pRP674 (CEN LEU2 rpb6Q100R)* | [8] |
| Y25602 | *MATa/MATα his3-Δ1/his3-Δ1 leu2-Δ0/leu2-Δ0 lys2-Δ0/LYS2 MET15/met15-Δ0 ura3-Δ0/ura3-Δ0 rpb6Δ::kanMX4/RPB6* | Euroscarf |
| YFN183 | *MATα his3-Δ1 leu2-Δ0 lys2-Δ0 met15-Δ0 ura3-Δ0 rpb6Δ::KanMX4 + pFL44L-RPB6 (2µm URA RPB6)* | This work |
| YFN637 | *MATα his3-Δ1 leu2-Δ0 lys2-Δ0 met15-Δ0 ura3-Δ0 rpb6Δ::KanMX4 + pRP674 (LEU2 CEN rpb6Q100R)* | This work |
| YFN629 | *MATα his3-Δ1 leu2-Δ0 lys2-Δ0 met15-Δ0 ura3-Δ0 rtr1Δ::kanMX4::HIS3 rpb6Δ::KanMX4 + pFL44L-RPB6 (2µm URA RPB6)* | This work |
| YFN634 | *MATα his3-Δ1 leu2-Δ0 lys2-Δ0 met15-Δ0 ura3-Δ0 rtr1Δ::kanMX4::HIS3 rpb6Δ::KanMX4 + pRP674 (CEN LEU2 rpb6Q100R)* | This work |
| YFN517 | *MATα ade2-1 his3-Δ200 leu2 lys2 trp1-Δ63 ura3-52 rpb7Δ::LEU2 pGEN-RPB7 (2µm TRP1 RPB7)* | [9] |
| yBF32 | *MATα ade2-1 his3-Δ200 leu2 lys2 trp1-Δ63 ura3-52 rpb7Δ::LEU2 + pGEN-rpb7-ΔC3 (2µm TRP1 rpb7-ΔC3)* | [9] |
| YFN632 | *MATα ade2-1 his3-Δ200 leu2 lys2 trp1-Δ63 ura3-52 rpb7Δ::LEU2 pGEN-RPB7 (2µm TRP1 RPB7) rtr1Δ::KanMX4* | This work |
| YFN633 | *MATα ade2-1 his3-Δ200 leu2 lys2 trp1-Δ63 ura3-52 rpb7Δ::LEU2 + pGEN-rpb7-ΔC3 (2µm TRP1 rpb7-ΔC3) rtr1Δ::KanMX4* | This work |

**Supplementary Table S2. Plasmids used**

| **Name** | **Yeast markers and ORI** | **Origin** |
|---|---|---|
| M4754 | *KanMX::HIS3* disruptor converter | [10] |
| pASZ11-*RPB5* | *ORI (CEN) ADE2* | [6] |
| pCM189 | ORI (CEN) *URA3* | [11] |
| pCM189-*BUD27* | ORI (CEN) *URA3* | [3] |
| pCM189-*RPB4* | ORI (CEN) *URA3* | [12] |
| pCM189-*RPB6* | ORI (CEN) *URA3* | This work |
| pFL44L | ORI (2µm) *URA3* | [13] |
| pFL44L-*RTR1* | ORI (2µm) *URA3* | This work |
| pFL44L-*RPB5* | ORI (2µm) *URA3* | [14] |
| *pFL44L-RPB6* | ORI (2µm) *URA3* | [4] |
| pGEN | ORI (2µm) *TRP1* | [15] |
| pGEN-*RPB7* | ORI (2µm) *TRP1* | [16] |
| pRS313-GFP-*RPB4* | ORI (CEN) *HIS3* | [17] |

**Supplementary Table S3. Oligonucleotides used**

| **Gene (or DNA region)** | **Primer** | **Sequence** |
|---|---|---|
| Intergenic region (chromosome V) | IntergChrV-F | TGTTCCTTTAAGAGGTGATGGTGAT |
| | IntergChrV-R | GTGCGCAGTACTTGTGAAAACC |
| *18S rDNA* | 18S-501 | CATGGCCGTTCTTAGTTGGT |
| | 18S-301 | ATTGCCTCAAACTTCCATCG |
| *ACT1* | *ACT1*-501 | GCCTTCTACGTTTCCATCCA |
| | *ACT1*-301 | GGCCAAATCGATTCTCAAAA |
| *GAL1* | *GAL1*-501 | TGGTGTTAACAATGGCGGTA |
| | *GAL1*-301 | GGGCGGTTTCAAACTTGTTA |
| *GAL10* | *GAL10*-501 | ACGGAGATTATGGTGCGTTC |
| | *GAL10*-301 | GGATTTTTGGGGCCTAAGAC |
| *HHT1* | *HHT1*-501 | TGCCTTTCCAAAGATTGGTC |
| | *HHT1*-301 | TTGGATAGTGACACGCTTGG |
| *MTG1* (-143/+61 pb) | *MTG1*-503 | TCAAAGAACACGGACCATCA |
| | *MTG1*-303 | TTGGTGTGAAGGATGAAACG |
| *MTG1* (+797/+930 pb) | *MTG1*-502 | TGCAAAATTTGAACGATGGA |
| | *MTG1*-301 | CCACTCGATAGCCGTTGATT |
| *PMA1* (-330/-234 pb) | *PMA1p*-503 | AAAGGCCAAATATTGTATTATTTTCAA |
| | *PMA1p*-302 | TTGGTGTTATAGGAAAGAAAGAGAAA |
| *PMA1* (+9/+116 pb) | *PMA1-5'*-501 | ACATCATCCTCTTCATCATCCTC |
| | *PMA1-5'*-301 | TCAGAAGATTCAGATGCAGCG |
| *PMA1* (+1367/+1532 pb) | *PMA1*-504 | GAAGGTTACTGCCGTTGTCG |
| | *PMA1*-304 | CGGAACCCTCTAGAAGCCAA |
| *PMA1* (+2551/+2757 pb) | *PMA1*-6 (forw) | ATATTGTTACTGTCGTCCGTGTCTGGAT |
| | *PMA1*-6 (rev) | ATTAGGTTTCCTTTTCGTGTTGAGTAGA |

| | | |
|---|---|---|
| *PYK1* (-37/+52 pb) | *PYK1p*-502 | ACAAGACACCAATCAAAACAAA |
| | *PYK1p*-301 | AGTCAGAACCAGCAACAACG |
| *PYK1* (+254/+352 pb) | *PYK1*-504 | CCAAGGGTCCAGAAATCAGA |
| | *PYK1*-304 | CGTACTTGTCATCGGTGGTG |
| *PYK1* (+769/+950 pb) | *PYK1*-503 | CTGACGGTGTTATGGTTGCC |
| | *PYK1*-303 | TCGGAAACTTCAGCTCTGGT |
| *PYK1* (+1039/+1288pb) | *PYK1*-4 (forw) | CTATGGCTGAAACCGCTGTCATTG |
| | *PYK1*-4 (rev) | CAGCTCTTGGGCATCTGGTAAC |
| *RPB4* | *RPB4*-502 | TGCTTGCAATGGTTCAGAAG |
| | *RPB4*-301 | CCATTTTTGGTCGAATTTTG |
| *RTR1* | *RTR1*-501 | TCCGATATTTTTGTCCGAAATAGG |
| | *RTR1*-301 | GACTCAAAGTGAATTATAGCAAAG |
| *URA2* (-123/+63 pb) | *URA2p*-501 | ATATCGGCATCTGGCTTGAA |
| | *URA2p*-301 | CAGACGGTCACCCGTAGATT |
| *URA2* (+618/+746 pb) | *URA2*-503 | GTACGTTCCTCCAGCAGACA |
| | *URA2*-303 | ACACCCCTTTTGATAAAACAACG |
| *URA2* (+2988/+3213 pb) | *URA2*-501 | CGAATTTGATTGGTGTGCTG |
| | *URA2*-301 | GGCGATGTTGTTGGAAGTTT |
| *URA2* (+6060/+6448 pb) | *URA2*-502 | CGCCGCTAAATATTCTCCTG |
| | *URA2*-302 | CAATTCCGGAGGAGAAACAA |
| 5'-Cy3 labelled probe | oligodTCy3 | TTTTTTTTTTTTTTTTTTTTTTTTTTTTTTTTTT |

The primers used were as follows (all 5’-3’)
